# Introgression from the wild relative *Manihot glaziovii* on cassava (*M. esculenta*) chromosome 1 exhibits segregation distortion and no direct effect on dry matter

**DOI:** 10.64898/2026.02.20.707074

**Authors:** Seren S. Villwock, Ismail Y. Rabbi, Andrew Smith Ikpan, Kayode Ogunpaimo, Kehinde Nafiu, Siraj Ismail Kayondo, Marnin Wolfe, Jean-Luc Jannink

## Abstract

The cassava (*Manihot esculenta)* genome has two large introgressions from its wild relative *M. glaziovii* on chromosomes 1 and 4 that originate from historical hybridization efforts. The 10 Mbp chromosome 1 introgression has been increasing in frequency in African breeding populations due to its statistical association with higher dry matter content and root number. However, the region also exhibits suppressed recombination, hindering breeders’ ability to combine favorable *glaziovii* alleles with the cultivated *esculenta* background. Since homozygous introgressed lines are rarely selected for advanced trials, dominance effects have not been well-characterized. To analyze the effects of the introgression with higher resolution, we generated a population of over 5000 seedlings from crosses between heterozygous introgressed parents and screened for recombinants using ten KASP markers tagging *glaziovii-*specific alleles. An optimized subset of 453 lines was then selected and evaluated over two years for yield and vigor traits. Unlike previous studies, composite interval mapping and mixed linear models showed no significant associations between *glaziovii* alleles and dry matter content or root number. Small, opposing effects on clonal vigor were observed at different ends of the introgression. The region showed significant segregation distortion and enrichment of putative deleterious alleles. Genome alignment of *M. esculenta* and *M. glaziovii* assemblies did not show any major structural variants in the introgression region, suggesting that suppressed recombination is likely driven by sequence-level divergence rather than structural rearrangements. These results indicate that the *glaziovii* introgression does not directly contribute to dry matter, supporting the need for recombination and purging of the *glaziovii* introgression to aid cassava improvement.

**Plain language summary:** A large chromosome segment from a wild relative of cassava is an important structural aspect in the cassava genome. Since the chromosome segment tends to be inherited as one block, its effects on cassava traits were not well resolved. Through genetic mapping at higher resolution, we identified that the wild segment impacts early vigor and does not appear to impact dry yield, as was previously thought. While there are no major structural differences between the wild and cultivated chromosome segments, their overall divergence seems to suppress the wild chromosome segment from pairing with the cultivated chromosome segment during reproduction. In the apparent absence of any major benefits from the wild segment, removing it from the breeding population may be beneficial.

**Core ideas:**

- A set of *glaziovii* allele-specific markers were designed to track the chromosome 1 introgression haplotype.
- Segregation distortion suggests the presence of recessive deleterious or lethal alleles in the introgression.
- Increased recombination is needed to purge deleterious alleles enriched in introgression region.
- The *glaziovii* introgression was associated with slightly lower vigor rating and stem diameter.
- The effects of the previously-identified *glaziovii* DM QTL were not detected in this population.

## 1 INTRODUCTION

Wild relatives are an important source of genetic diversity in plant breeding, particularly for traits that have not historically been prioritized for selection. Making interspecific crosses with wild relatives can introduce new or lost alleles into breeding populations but can also carry along deleterious alleles via linkage drag. Through backcrossing and recombination, breeders aim to isolate the beneficial wild alleles while recovering as much as possible of the elite background. This process can be challenged when beneficial and deleterious alleles are closely linked on the introgressed haplotype. It is further complicated by suppressed meiotic recombination rates, which have been associated with introgression regions in interspecific hybrids due to sequence divergence or structural variants that can inhibit homologous pairing (Lawrence et al., 2017; Delame et al., 2019; Rowan et al., 2019).

In the 1930s and 40s, cassava (*Manihot esculenta*) breeders at the Amani Research Station in present-day Tanzania made interspecific crosses to introgress disease resistance from wild relatives. They sought to breed new varieties with resistance to cassava mosaic disease (CMD) and cassava brown streak disease (CBSD), two viral diseases that threaten the crop yields and sustenance of millions of smallholder farmers in East Africa (Hillocks & Jennings, 2003). Progeny descending from crosses between *M. esculenta* and its tree-like wild relative *M. glaziovii* became important founders for modern breeding programs (Hillocks & Jennings, 2003). Today, there are two large (10-20 Mb) *glaziovii* introgression segments on cassava chromosomes 1 and 4, a remnant from those hybridizations (Bredeson et al., 2016). Wolfe et al. (2019) found that these *glaziovii* introgressions are common in the African breeding populations and have increased in frequency over cycles of genomic selection.

Interestingly, neither *glaziovii* introgression appears to be associated with cassava mosaic disease resistance, the original motivation for making the crosses, but they have been associated with favorable effects on other traits (Wolfe et al., 2019). The chromosome 1 *glaziovii* introgression (C1GI) is of particular interest because it harbors important quantitative trait loci (QTL) for root number and dry matter (DM) content, a key breeding focus for improved yield and textural qualities that determines varietal adoption (Tahirou et al., 2015). *Glaziovii* alleles contributed 4.5% and 16.1% of the heritability of these traits, respectively, in Wolfe et al. (2019). However, the introgressions are also enriched for putative deleterious mutations, suggesting that these regions are affected by linkage drag and would benefit from recombination to purge unnecessary genetic load (Ramu et al., 2017b; Wolfe et al., 2019). Being able to fix the *glaziovii* QTL in a homozygous state would capture the full additive benefit of the favorable allele(s) and allow breeders to maintain them more easily in the population.

Unfortunately, both introgression regions have significantly suppressed recombination rates (Chan et al., 2022). The 10Mbp C1GI has a genetic length of about 35cM in non-introgressed lines, while homozygous introgressed lines have a genetic distance of about 10cM across the same region (Chan et al., 2022). Suppressed recombination limits the resolution of mapping studies and makes it difficult to break or fix linkages in this region. Notably, there is hypothesized to be a repulsion linkage between a favorable dry matter allele and unfavorable carotenoid content allele, which may drive the negative genetic correlation between provitamin A and dry matter contents (Rabbi et al., 2017; Gemenet et al., 2020).

Another challenge that the C1GI poses to cassava breeding is that it is typically maintained in a heterozygous state, so the QTL continue to segregate in crosses. Although there is a high level of heterozygosity across the cassava genome in general, the C1GI region is particularly important because it harbors the strongest linkage disequilibrium (LD) block in the cassava genome and contains the dry matter QTL with the largest effect size identified to date (Rabbi et al., 2017). Homozygous introgressed lines are not well-observed because they are almost completely excluded from advanced trials (Wolfe et al., 2019).

The exclusion of homozygotes may be due to the enrichment of putative deleterious mutations on the *glaziovii* haplotype (Wolfe et al., 2019). It is likely that the C1GI haplotype descended from the same *esculenta* x *glaziovii* hybrid ancestor and thus has low genetic diversity compared to the *esculenta* haplotypes and is less tolerant of homozygosity. Homozygous lines may also be lost early on due to poor germination, vigor, or survival. Before being genotyped or advanced to clonal evaluation trials (CETs) for yield evaluations, seedlings are typically screened for disease resistance, desirable plant architecture, and aboveground vigor, which are assessed visually on a categorical scale. We hypothesized that the C1GI introgression carries deleterious load that negatively impacts plant vigor, a target of early rogueing in seedling nurseries. The distribution of putative deleterious alleles throughout the introgression region was examined to assess whether particular C1GI segments could be targeted for purging.

While the C1GI has an important influence on dry matter content, a major trait prioritized in West African breeding programs, the extensive LD in the region limits the resolution of QTL analysis. Wolfe et al. (2019) identified *glaziovii* SNPs significantly associated with dry matter content over a 6 Mbp region, from about 24-30 Mbp. Questions remain about which segment of the introgression carries the QTL, its distance from the root number QTL, if there are multiple distinct loci affecting the trait, and if the QTL is separable from other effects of interest, including a non-favorable effect on carotenoid content associated with the same region (Rabbi et al., 2017). Recombining the introgression will increase the resolution for QTL analysis and could generate a more favorable haplotype that isolates the beneficial QTL to a smaller region that would be less impacted by linkage drag.

The overall approach of this project was to generate a diverse population that contained more recombination in the C1GI region to map both its advantageous and deleterious effects with higher resolution. A large population of seedlings was screened to identify a subset of lines that would maximize the statistical power to isolate effects of introgression segments. We aimed to determine optimum recombination breakpoints that can be identified with genetic markers to select the most favorable segment(s) of the C1GI while purging genetic load. Overall, these analyses provide an interesting case study into the long-term effects of interspecific hybridizations in breeding an outcrossing crop with inbreeding depression.

## 2 METHODS

### Cross selection and seedling nursery

The C1GI status of previously genotyped accessions in the 2020 crossing nursery in Ubiaja, Nigeria was examined using the introgression diagnostic markers (IDMs) identified by Wolfe et al. (2019). To generate a mapping population with as many recombinations in the C1GI region as possible, accessions that were heterozygous for all or part of the C1GI were selected as parents. These 41 parents were clustered into three groups based on genetic distances derived from pairwise kinship estimates and clustered with the *hclust()* function in R (R Core Team, 2021). The groups were intercrossed, and open-pollinated seeds from these parents were also collected. A total of 5062 seedlings from 272 families were planted in a seedling nursery (SN) at IITA Ibadan, Nigeria in early April 2021. The generated population was therefore analogous to an F2 intercross population with respect to the C1GI haplotype, though with 41 parents and therefore more diversity in the rest of the genome. Germinated seedlings were transplanted 0.5m apart in rows 1m apart. Field trial management was conducted following standard practices (Abass et al., 2014).

### Marker design and genotyping

To screen the seedling nursery population for C1GI recombinant accessions, a panel of ten low-cost ‘Kompetitive’ Allele-Specific Primer (KASP) markers were designed to track the C1GI haplotype. These SNP markers were adapted from the Wolfe et al. (2019) *M. glaziovii* IDMs but validated across a larger reference panel to ensure consistent amplification across a diverse population. *M. glaziovii*-specific alleles across the C1GI were identified using a reference panel of Cassava HapMapII (Ramu et al., 2017b) accessions containing seven *M. glaziovii* individuals, 25 individuals heterozygous for the full C1GI, and 68 non-introgressed *M. esculenta* individuals, as determined by the previously identified IDMs (Wolfe et al., 2019). Then, genotyping-by-sequencing SNPs in this panel were filtered with *bcftools* using the following criteria: 1) reference allele fixed within the non-introgressed *esculenta* panel, alternate allele fixed within the *M. glaziovii* panel, and one copy of the reference allele in the heterozygous introgressed accessions; 2) SNP called in more than 90% of the population; 3) average read depth per site in the surrounding 100 bp region between 2.2-3.5; 4) minimum and maximum read depths at sites in the surrounding region 0.86 and 3.50, respectively; 5) no indels in the surrounding region; and 6) two or fewer other SNPs in the surrounding 100bp region.

The 38 candidate SNPs and their surrounding sequences were then tested on the 41 mapping population parents, three *M. glaziovii* trees, and an *M. esculenta x glaziovii* hybrid growing onsite at IITA Ibadan. Freeze-dried leaf tissue samples were sent to Intertek (Intertek Australia Lab, Parkside South Australia) for DNA extraction and KASP marker genotyping. A final set of ten markers were selected that showed good amplification, tight clustering, and even spacing across the C1GI. We validated the markers by evaluating whether the wild *M. glaziovii* accessions, hybrid *M. glaziovii* x *esculenta* accessions, two non-introgressed checks (TME419 and TMS18F1279P0006) clustered with their expected genotype classes. A random subset of 3006 progeny from the seedling nursery were then genotyped with the KASP marker panel to determine their introgression status.

The subset of 453 clones advanced to the CET (described below) were also genotyped genome-wide with DArTSeqLD (Diversity Array Technologies, Canberra, Australia). The 12,981 DArTSeqLD SNPs were imputed to higher density with Beagle 5.0 (Browning & Browning, 2016) using the R package genomicMateSelectR version 0.2.0 (Wolfe, 2022) with a reference panel of 21,856 accessions available on Cassavabase (https://cassavabase.org/ftp/marnin_datasets/nextgenImputation2019/). Following imputation, SNPs were filtered with the following parameters: dosage *R*^2^ (DR2) > 0.75, Hardy-Weinberg equilibrium p-value (P_HWE) > 1e-20, and minor allele frequency (MAF) > 0.05 for a total of 58,830 biallelic SNPs.

### Optimizing a mapping population subset for replicated trials

To assess root yield in a clonal evaluation trial (CET) with space for 500 plots, a subset of accessions was selected to capture as much diversity and recombination in the introgression haplotypes as possible. The subset was selected using Federov’s exchange algorithm implemented by the *OptFederov()* function in the AlgDesign R package with the *D* criterion, which maximizes the determinant of the cross-product of the design matrix to capture as much information about the model parameters as possible (Wheeler, 2019). The additive and dominance effects of all ten KASP markers were included together in the model as follows:

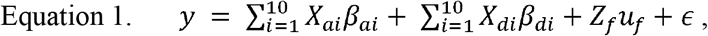

where *y* is the vector of phenotypic values; *X*_*ai*_ is the matrix of genotypes at marker *i* in {0,1,2} encoding of the dosage of the *M. glaziovii* allele; *β*_*ai*_ is the additive fixed effect of KASP marker *i*,; *X*_*di*_ is the matrix of genotypes at marker *i* in {-1,1,-1} encoding for the *M. esculenta* homozygote, heterozygote, and *M. glaziovii* homozygote, respectively; *β*_*di*_ is the dominance fixed effect of marker *i*; *Z*_*f*_ is the design matrix of families; *u*_*f*_ is the vector of family random effects assumed 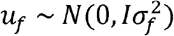; and *ϵ* is the vector of residual effects assumed 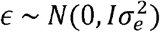.

The *OptFederov* algorithm was run ten times, given the parental genotypes to include by default and the genotypes of 2981 seedlings without any missing SNPs to select from. The subset of 400 lines with the highest *D* value at 0.057 was then selected and propagated to a CET, with each genotype planted in one plot of five plants. An additional 4 check lines (IITA-TMS-IBA000070, IITA-TMS-IBA070593, TMEB419, and TMS13F1343P0022) and 39 plants identified as promising clones for breeding purposes by the breeders’ phenotypic evaluation and were also planted for a total of 453 accessions in the CET.

### Testing segregation ratios

The progeny of two heterozygous parents at each C1GI marker were tested for deviation from the expected 1:2:1 segregation ratio using chi-square goodness of fit tests for the genotype counts at each KASP marker. To evaluate the possible impact of pollen contamination on deviations from expected genotype frequencies, the parentage assignment algorithm AlphaAssign (Whalen et al., 2019) was used to estimate the error rate of pedigree records using the genome-wide SNPs of the 453 CET individuals and accessions originally present in the crossing nursery. The seed parent listed on the pedigree was input into AlphaAssign with a list of all clones present in the crossing nursery as potential pollen parents. Pollen contamination rate was estimated based on how often the AlphaAssign algorithm identified a different pollen parent than the one recorded in the pedigree.

To determine if the pollen contamination rate would be large enough to explain the observed deviation from the expected segregation ratios, a simulation was conducted to model changes in genotype frequencies when the pollen parent was randomly selected from the crossing nursery population at the observed pedigree error rate. A total of 812 F_2_ progeny (the largest number of progeny with both parents heterozygous at a given C1GI marker; Table 2) were simulated with ten marker genotypes drawn from the probabilities expected if all parents were heterozygous and there was no segregation distortion ({P_0_ = 0.25, P_1_ = 0.5, and P_2_ = 0.25}). Lines representing the pollen contamination errors, based on the rate estimated with AlphaAssign, were randomly selected. New genotypes were assigned to these lines based on the expected genotype frequencies when a heterozygote is pollenated with probability (1-*q*) of receiving an *esculenta* allele and probability *q* of receiving a *glaziovii* allele ({P_0_ = 0.5*(1-*q*), P_1_ = 0.5*(1-*q*), and P_2_ = 0.5\**q*}, where *q* is the frequency of the *M. glaziovii* allele in the parental population (assuming all accessions were equally likely to be a contaminating pollen parent). The resulting population was then tested for deviation from a 1:2:1 ratio for each marker with chi-square tests. This simulation was repeated 1000 times and the number of times the chi square test for each marker was significant (p < 0.05) were counted as false positives. The significance threshold was then adjusted to empirically restrict the false-positive rate to 5% accounting for the pedigree error rate.

### Phenotyping

The population was grown in five total field trials across four years and two locations. The trial stages included the seedling nursery (SN; unreplicated plants grown from seed, 5062 genotypes of which 3006 had KASP marker data, planted by family), 2 clonal evaluation trials (CET; 1 plot of 5 plants per genotype, 453 genotypes, augmented design), and 2 preliminary yield trials (PYT; 2 plots of 10 plants per genotype, 50 genotypes, randomized complete block design).

Agronomic traits including cassava mosaic disease (CMD) severity, root number, root size, root weight, dry matter content, plant architecture, branching type, sprouting ability, and chromameter values were measured according to standard practices (Fukuda et al., 2010) (Table 2). Agronomic traits were measured per-plant in the SN (Ibadan, 2020), per-plot of five plants in the CETs (Ibadan 2021 and Ikenne 2022) and per-plot of ten plants replicated twice in the PYTs (Ibadan 2023 and Ikenne 2023). Correlations between traits were tested with Kendall’s rank correlation (τ) since the yield data were right-skewed.

In the CGIAR cassava trait ontology, plant vigor is scored on a three-point ordinal scale (Fernandez-Pozo et al., 2015). To capture more quantitative variation in vigor, plant height and stem diameter were also measured as traits that contribute to the breeder’s visual assessment of overall aboveground vigor. Plant height was measured from the soil to the apical meristem with a meter stick. Stem diameter was measured with a caliper at soil level for the seedlings and at the base of the new stem for clonally propagated plants; in the case of multiple basal stems, the diameter of the largest stem was taken. Vigor traits were measured on individual plants in the SN at 4 months after planting (MAP) and on three plants per plot in the CETs at 6 and 8-9 MAP.

Vigor traits measured on plants grown from seed in the SN trial and grown from stem cuttings in the clonal trials were considered separate traits since they correspond to different biological processes and had such different sample sizes (3,006 seedlings vs. 453 clones). Therefore, seedling and clonal traits were modeled separately as described below. Associations with yield traits in the SN were not reported since single-plant measures of yield are generally considered unreliable.

### Genetic mapping

Associations between C1GI markers and all traits were analyzed with two approaches: mixed linear models (MLM) with individual markers and composite interval mapping (CIM) accounting for multiple markers (Zeng, 1994). First, linear models for SN vigor traits (unreplicated plant height and stem diameter measurements for the 3,006 progeny genotyped only with the 10 C1GI markers) were fit separately as follows:

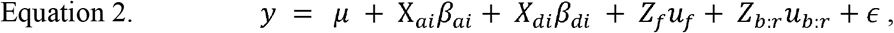

with variables as described in Eqn. 1, and where *μ* is the grand mean, *Z*_*b:r*_ is the design matrix for rows within field blocks, and *u*_*b:r*_ is the vector of row/block random effects assumed 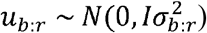. This model was fit for each C1GI marker *i* using the *lme4* R package v1.1-35.5 (Bates et al., 2015). The significance of marker additive and dominance effects was assessed by testing the null hypotheses that *β*_*a*_ =0 and *β*_*d*_ =0 with Wald tests. For multiple testing correction, the number of independent tests was estimated as described in Gao et al., (2008); since the first 5 eigenvalues explained > 99.5% of the variation in the *glaziovii* KASP marker correlation matrix, a significance threshold of *α* = 0.05 was established at p < 0.01.

Associations for all vigor and agronomic traits among CET-selected accessions (replicated measurements across environments for the 453 accessions genotyped with both whole-genome markers and the C1GI markers) were conducted in two steps. First, best linear unbiased predictions (BLUPs) of genotype effects on agronomic traits were calculated using the following model using the *ASReml-R* package v4.1.0.110 (The VSNi Team, 2023):

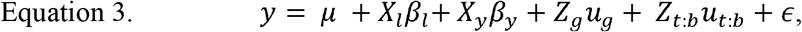

with variables as described in Eqn. 1 and 2, and where *X*_*l*_ is the design matrix for trial locations, *β*_*l*_ is the vector of fixed effects for locations, *X*_*y*_ is the design matrix for trial years, *β*_*y*_ is the vector of fixed effects for years, *Z*_*g*_ is the design matrix for genotypes, and *u*_*g*_ is the vector of genotype random effects assumed 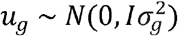 where 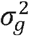 is the genetic variance, *Z*_*t:b*_ is the design matrix for blocks nested within trials, *u*_*t:b*_ is the vector of random effects for blocks within trials assumed 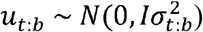, and *ϵ*is the vector of residual effects assumed *ϵ ∼ N(0, R)*, where 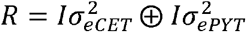 to account for heterogeneous error variance across trial types. For the vigor traits with replicated measurements (stem diameter and plant height each measured on three plants per plot at two timepoints, 4 MAP and 8/9 MAP), an additional fixed effect for timepoint and a random effect interaction between genotype and timepoint were added to the models.

The estimated breeding values (EBVs), *u*_*g*_, were extracted from these models and de-regressed by dividing by their prediction reliability 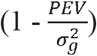 as described in Garrick et al., 2009. The broad-sense heritability (H^2^) of each trait was calculated as follows 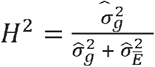, where 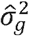 is the estimated genetic variance and 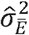 is the weighted average of residual error variance estimates, 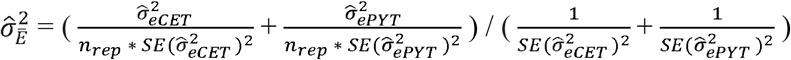, where *n*_*rep*_ is the total number of plot replicates across locations (2 for CETs and 4 for PYTs) (Holland et al., 2002; Schmidt et al., 2019). Then, the de-regressed EBVs were used for the MLM and CIM association analyses. For the MLM method, marker effects were estimated with the following model using the *ASReml-R* package v4.1.0.110 (The VSNi Team, 2023):

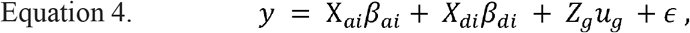

with variables as described above except *y* is now the vector of de-regressed EBVs and *u*_*g*_ is the vector of genotype random effects assumed 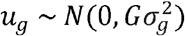, where *G* is a genomic relationship matrix calculated with *rrBLUP* package v4.6.3 (Endelman, 2011) using the DArtSeqLD imputed SNPs for chromosomes 2-18 (excluding chromosome 1 to separate C1GI effects from rest-of-genome effects). The significance of marker additive and dominance effects were tested with Wald tests as described above.

The CIM approach was then used to account for multiple marker effects simultaneously and distinguish whether there are multiple QTL across the C1GI. We used the *cim()* function in the R package *qtl* v1.7 (Broman et al., 2003) to scan the C1GI region using the Kosambi mapping function with assumed genotyping error rate of 0.0001, window size of 2 cM, up to 8 C1GI marker covariates, the first 10 principal components of the DArtSeqLD imputed genotype matrix from chromosomes 2-18 to account for population structure in the rest of the genome, the first 4 principal components of the non-introgression region of chromosome 1 as covariates to account for effects elsewhere on chromosome 1, and a previously published genetic map from Chan et al. (2022) uplifted to coordinates for cassava reference genome v7. An LOD (logarithm of the odds) threshold corresponding to *α* = 0.05 was determined with a permutation test of 1,000 permutations for each trait. QTL effect estimates were calculated with the *fitqtl()* function using the Haley-Knott regression method (Haley & Knott, 1992; Broman et al., 2003).

### Comparative genomics

One hypothesis for suppressed recombination in interspecific introgressions is that large structural variants (SVs) between haplotypes impedes homologous pairing (Li et al., 2023). While some structural variants can be inferred from patterns of read depth in short paired-end sequencing data, the detection ability is limited to certain types of SVs and typically has poor sensitivity and high false-positive rates (Sarwal et al. 2022). To look for C1GI structural variants that might have been missed by alignment to the cassava reference genome (which lacks the C1GI), we used a new haplotype-resolved *M. glaziovii* genome assembled with PacBio HiFi, Nanopore, and HiC sequencing by the Salk Institute Michael Lab (*Mgla*.*hap1*.*v1* and *Mgla*.*hap2*.*v1*, available at https://resources.michael.salk.edu/root/tools.html?tool=%2Fresources%2Fcassava_genomes%2Findex.html). The *M. glaziovii* haplotype assemblies were aligned to *M. esculenta* AM560-2 reference v7.1 (Bredeson et al., 2016) using AnchorWave v1.0.1 *proali* with maximum alignment parameters set to 1 for both reference and query (Song et al., 2022). Structural variants were called using SVIM-asm in diploid mode (Heller & Vingron, 2020).

A previous study by (Wolfe et al., 2019) found that the C1GI was enriched with putative deleterious alleles that were identified by Ramu et al. (2017) using SIFT (“Sorting Intolerant From Tolerant”) scores that predict the functional impact of mutations on proteins. Long et al. (2023) improved on the SIFT method of identifying deleterious alleles by including evolutionary conservation across the Euphorbiaceae family. To test the hypothesis that the C1GI carries a higher deleterious load, the proportion of putative deleterious SNPs to non-deleterious SNPs (as defined by Long et al., 2023) within the C1GI region was compared to that of an equal number of randomly selected SNPs from elsewhere in the genome across 1,000 permutations.

## 3 RESULTS

### KASP marker design and genotyping

The amplification of the ten *glaziovii* KASP markers on 3,006 tissue samples from a random subset of the SN and the parents is shown in Fig. 1. The number of non-callable genotypes ranged from 79 to 96 per marker, corresponding to an average success rate of 97.2%. The check individuals clustered as expected with the respective *esculenta*, heterozygous, and *glaziovii* clusters (Fig. 1). Adjacent marker pairs had between 11 and 458 individuals with a recombination event between them identified. An optimized subset of 400 individuals were selected for replication in the CET (Fig. 2). The marker genotypes were highly correlated in the seedling nursery (adjacent markers with *r =* 0.81-0.99), as was expected due to their physical proximity about 1 Mbp apart in a region with suppressed recombination. Correlations decreased slightly in the CET due to the selection of recombinants (Fig. 3).

**Figure 1.**
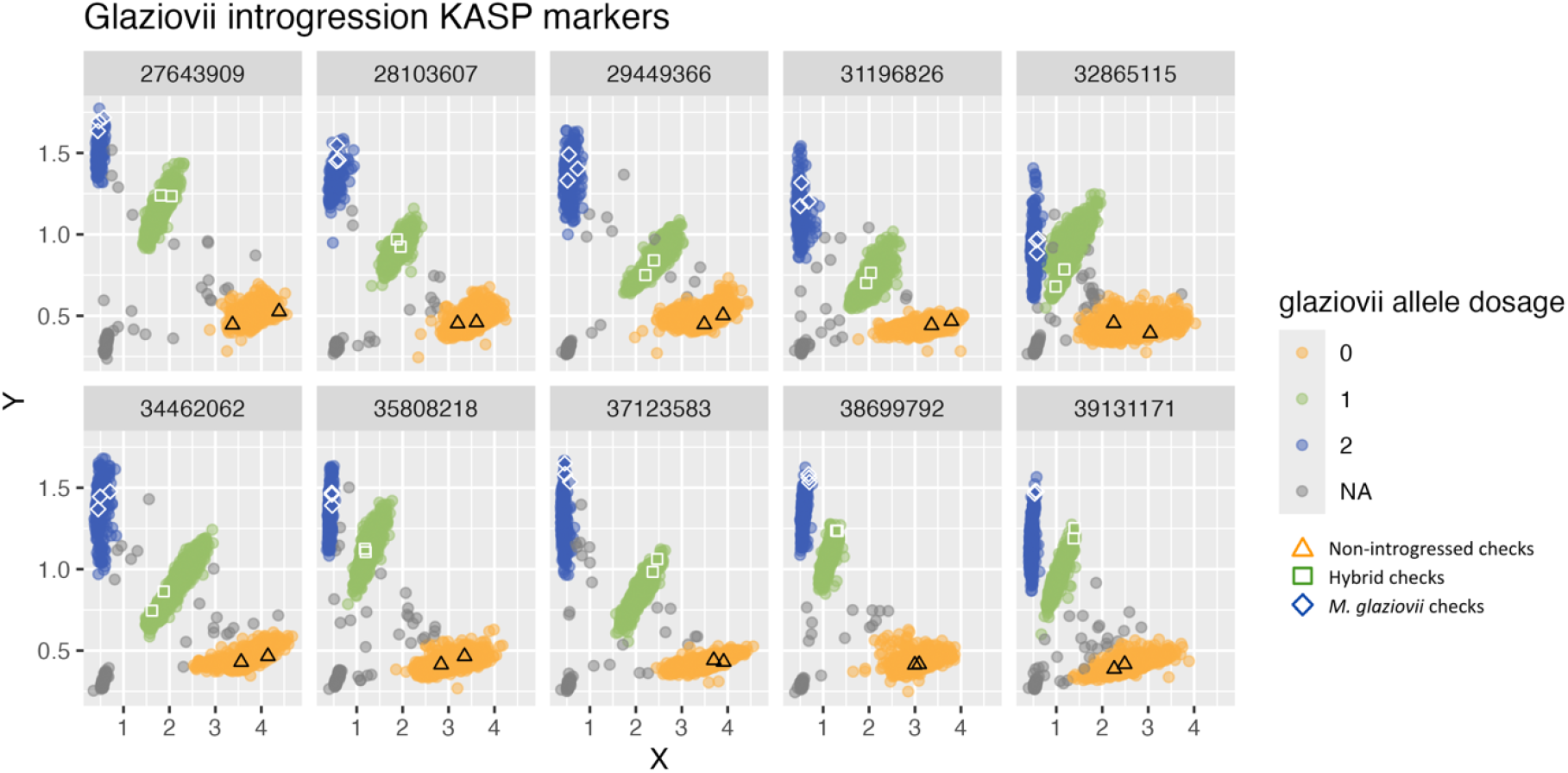
KASP marker panel tagging ten glaziovii-specific alleles across the introgression. The wild M. glaziovii, hybrid esculenta x glaziovii, and non-introgressed esculenta checks (TME419 and TMS18F1279P0006) are represented by blue diamonds, green squares, and orange triangles, respectively. The top of each panel gives the chromosome 1 marker position in base pairs. X and Y axes are fluorescence readings from KASP primer amplification of the two alleles. NAs represent individuals for which the marker genotype could not be called unambiguously.

**Figure 2.**
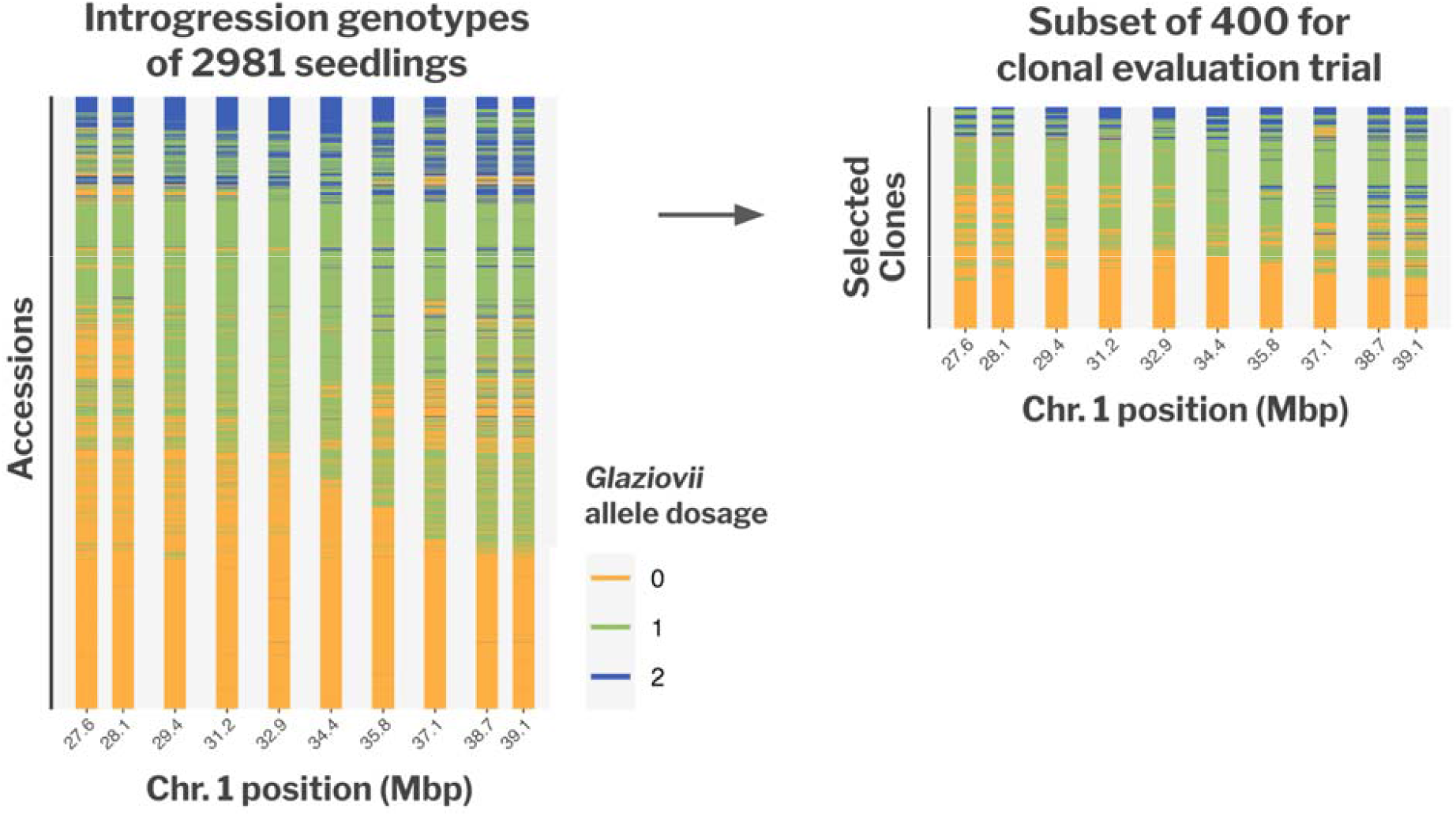
Selection of an optimum subset of seedlings to advance to a clonal evaluation trial for analysis of M. glaziovii introgression effects. The bars show glaziovii allele dosage across the ten C1GI KASP markers. The 2981 individuals in the seedling nursery without missing SNPs are shown at left and the selected seedlings and their parents are at right. The CET selection was optimized with Federov’s exchange algorithm to maximize information captured about marker and family effects.

**Figure 3.**
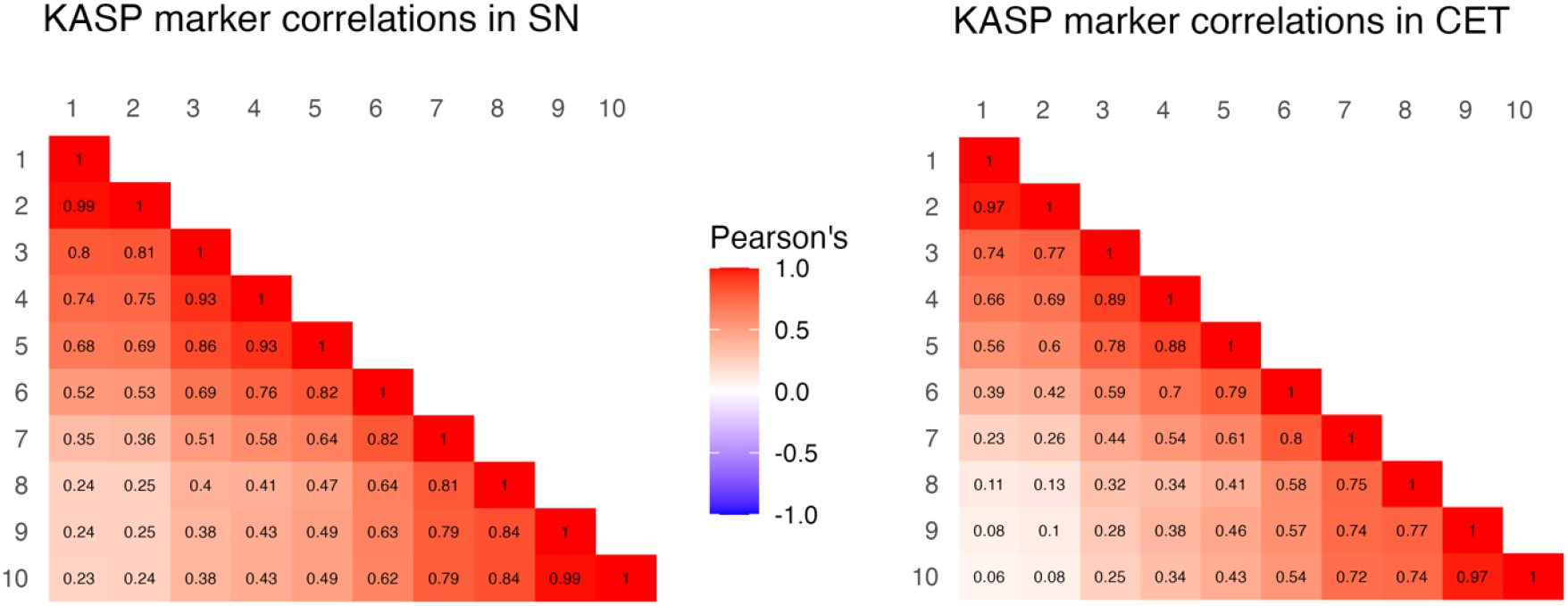
Pairwise correlations between C1GI KASP markers. The heatmap at left shows the Pearson’s correlation coefficients between markers among the seedling nursery population of N = 3,006, and at right are the correlations among the clonal evaluation trial of N = 453 after selection for recombinants.

Across all 10 markers, fewer homozygous introgressed progeny were observed than expected under a 1:2:1 Mendelian segregation ratio (Table 1). To test whether the observed segregation distortion could be an artifact of pollen contamination with a random pollen parent that may not have been heterozygous, pedigree errors were simulated to evaluate its impact on genotype frequencies. The parental assignment algorithm AlphaAssign identified a different pollen parent than the one recorded on the pedigree 16% of the time. Simulating this error rate in the population as a random pollen parent from the crossing nursery instead of the recorded heterozygous pollen parent resulted in a significant chi-square test, a false positive for segregation distortion, 39% of the time. To constrain the false positive rate at 5%, we set an empirical significance threshold at p < 0.0007. All of the C1GI markers still exhibited significant segregation distortion at this stricter significance threshold, indicating that pedigree error does not fully explain the observed segregation distortion (Table 1).

**Table 1.**
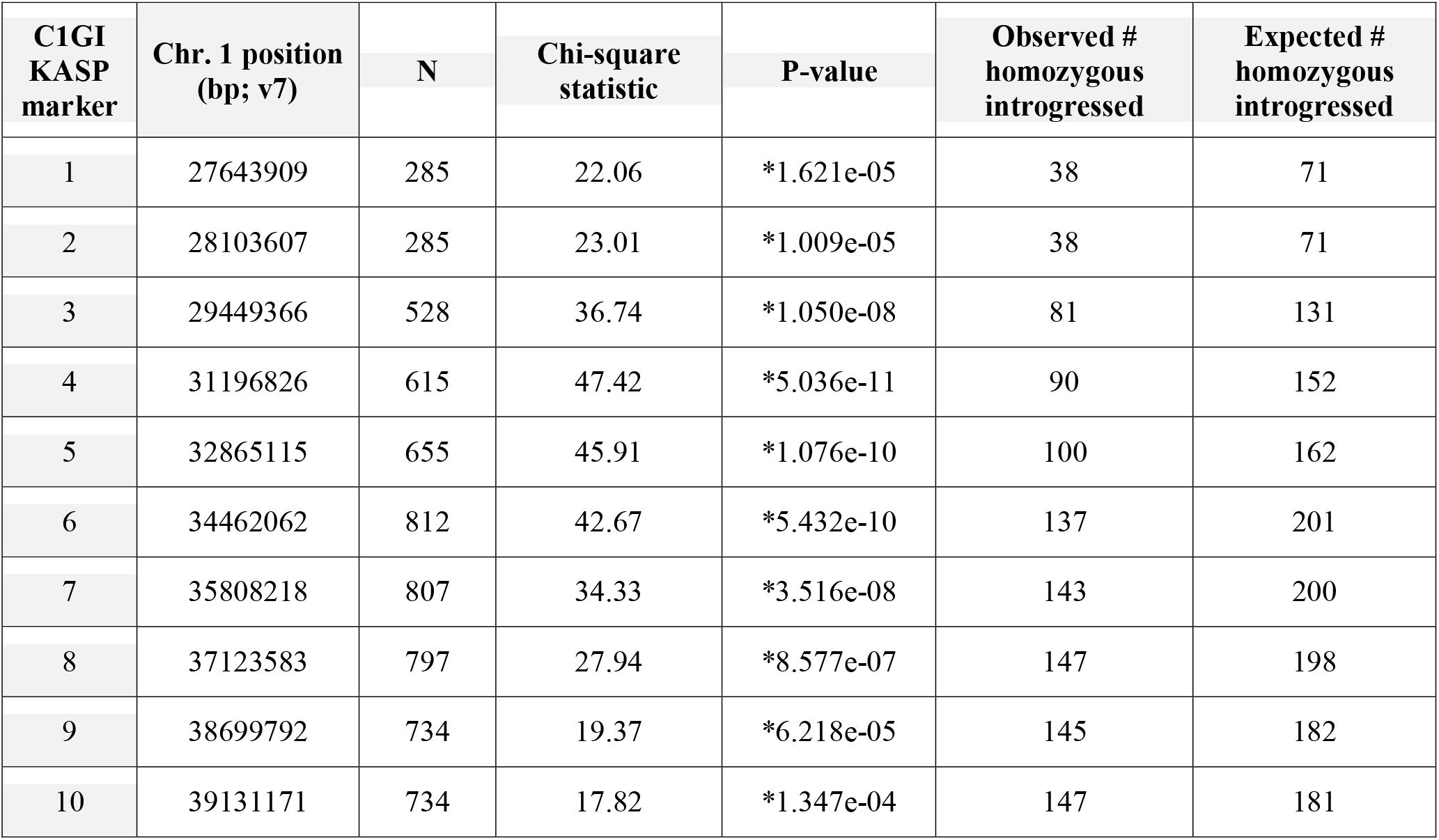
Chi-square tests for deviation from the expected 1:2:1 segregation ratio among progeny in the seedling nursery. N indicates the number of progeny included in a given marker test, for which both parents were heterozygous at that marker. Asterisks indicate p-values significant at the empirically-derived threshold of p < 0.0007 accounting for the estimated pedigree error rate.

| C1GI KASP marker | Chr. 1 position (bp; v7) | N | Chi-square statistic | P-value | Observed # homozygous introgressed | Expected # homozygous introgressed |
| --- | --- | --- | --- | --- | --- | --- |
| 1 | 27643909 | 285 | 22.06 | *1.621e-05 | 38 | 71 |
| 2 | 28103607 | 285 | 23.01 | *1.009e-05 | 38 | 71 |
| 3 | 29449366 | 528 | 36.74 | *1.050e-08 | 81 | 131 |
| 4 | 31196826 | 615 | 47.42 | *5.036e-11 | 90 | 152 |
| 5 | 32865115 | 655 | 45.91 | *1.076e-10 | 100 | 162 |
| 6 | 34462062 | 812 | 42.67 | *5.432e-10 | 137 | 201 |
| 7 | 35808218 | 807 | 34.33 | *3.516e-08 | 143 | 200 |
| 8 | 37123583 | 797 | 27.94 | *8.577e-07 | 147 | 198 |
| 9 | 38699792 | 734 | 19.37 | *6.218e-05 | 145 | 182 |
| 10 | 39131171 | 734 | 17.82 | *1.347e-04 | 147 | 181 |

**Table 2.**
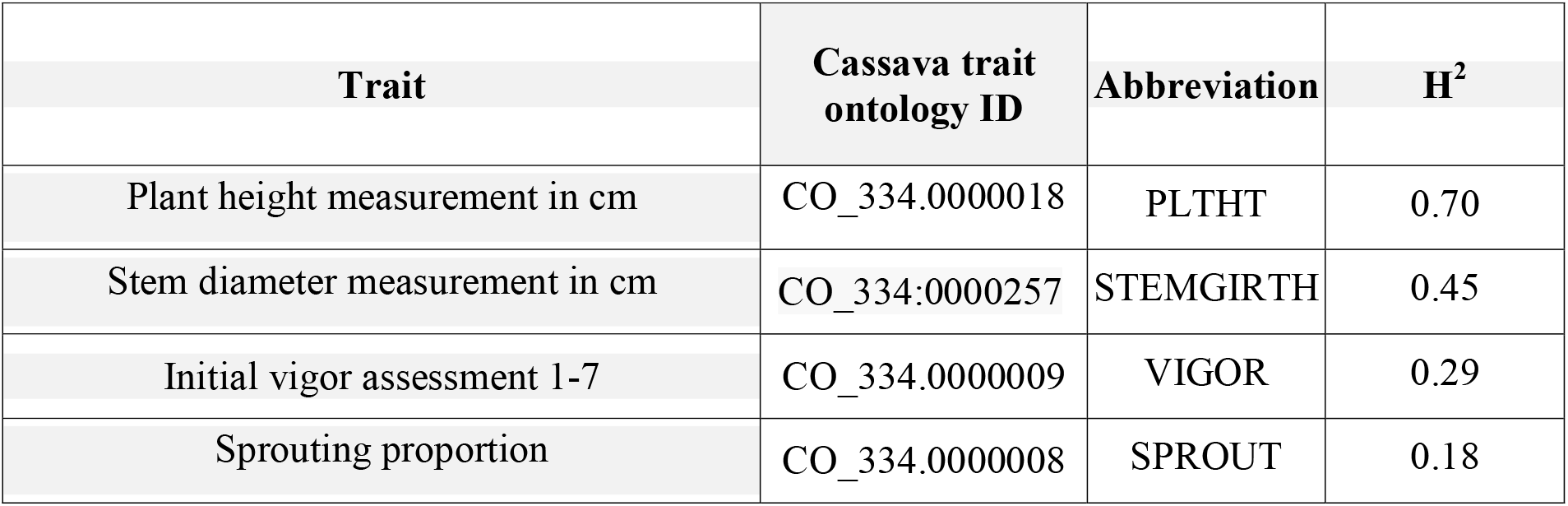
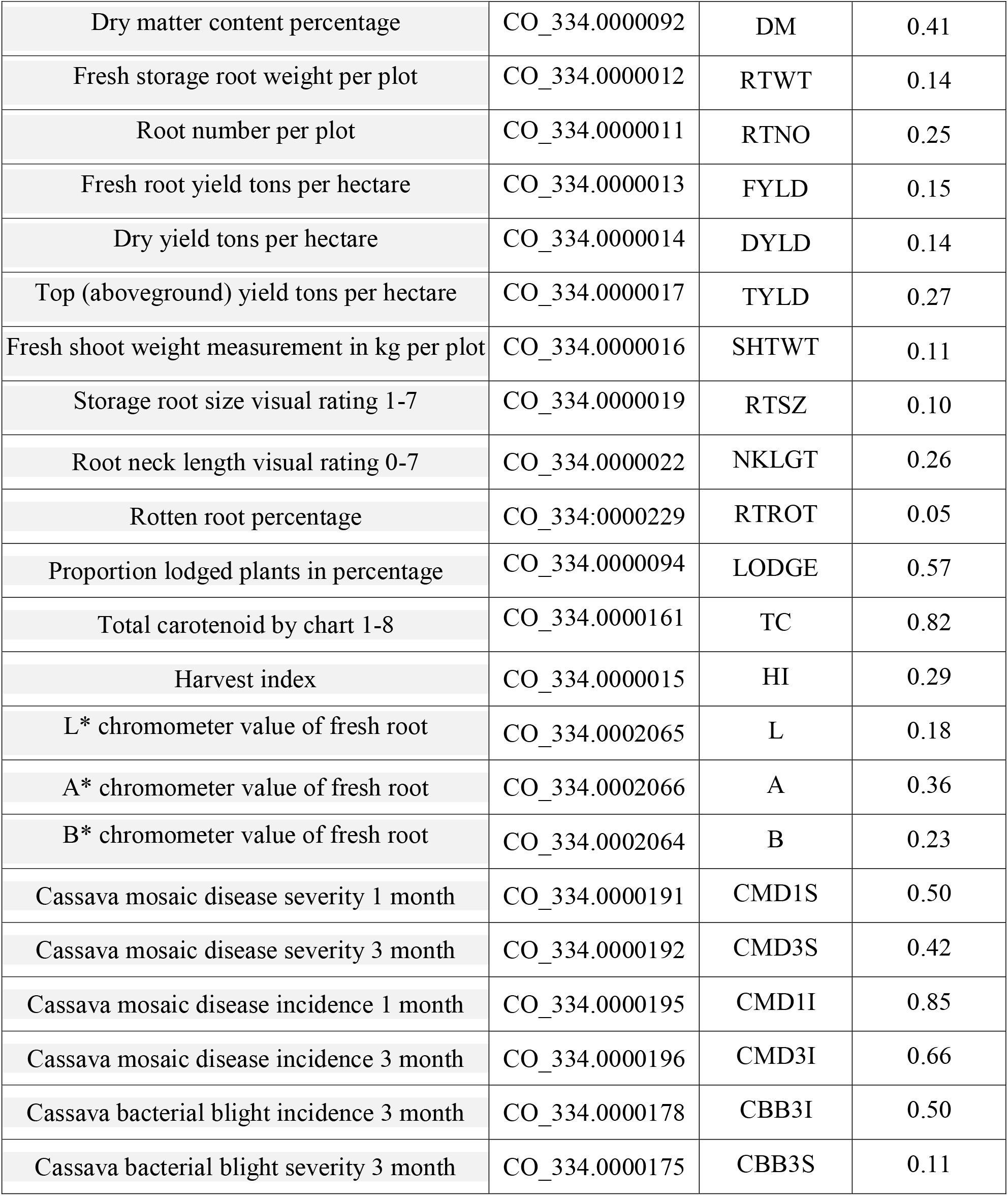
Trait broad-sense heritabilities. Heritabilities were estimated using variance components extracted from the linear model shown in Eqn. 3.

### Vigor trait association analyses

Vigor traits in the seedling and clonal trials were analyzed separately. In the SN trial, stem diameter at 4 MAP was significantly correlated with root weight (τ = 0.509, p < 2.2x10^-16^), root number (τ = 0.348, p = 7.1x10^-13^), and root size rating (τ = 0.249, p = 4.3x10^-6^). Both quantitative measurements of vigor, stem diameter and plant height, were significantly correlated with the breeder’s categorical rating of vigor (p = 4.2x10^-12^; p = 1.1x10^-14^ respectively).

With the MLM association method, C1GI markers 3 and 5-10 had positive dominance effects on seedling stem diameter, though the effect estimates were small (∼ 0.3 mm) (Fig. 4A). The CIM method distinguished two separate QTL associated with seedling stem diameter, with significant LOD peaks at marker 6 (estimated additive effect = -0.79 mm; dominance effect = 0.57 mm) and marker 8 (estimated additive effect = 0.43 mm; dominance effect = 0.40 mm) (Fig. 5A). There were no significant associations for SN plant height using either the MLM or CIM methods (Fig. 4B, Fig. 5B).

**Figure 4.**
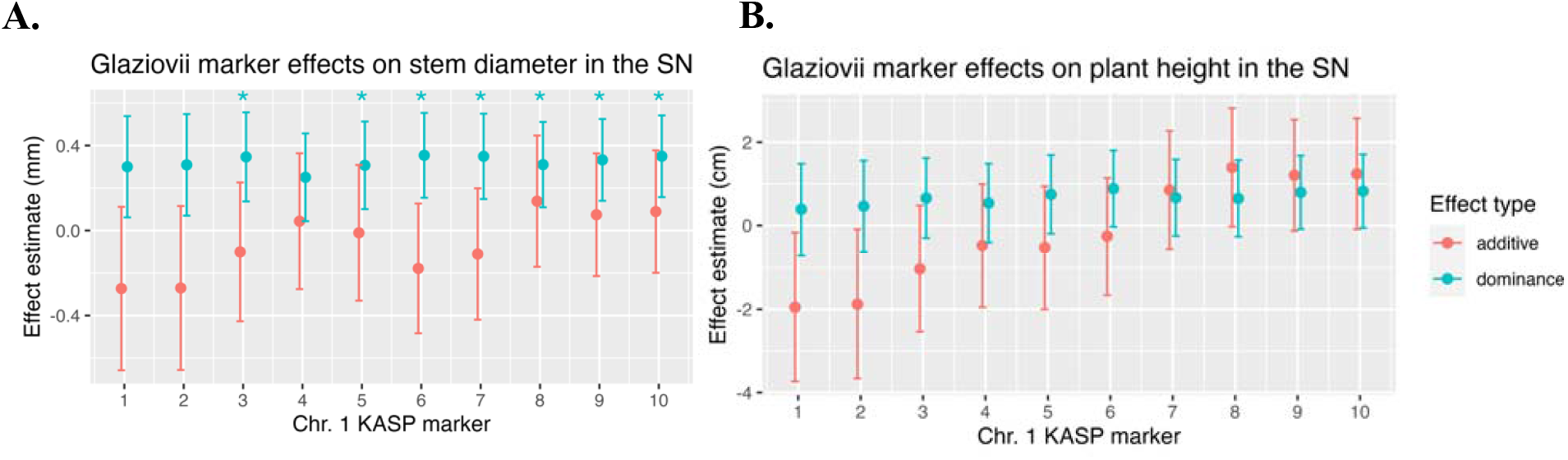
Estimates of additive (red) and dominance (blue) effects of M. glaziovii introgression markers on seedling vigor traits: A) stem diameter (left) and B) plant height. Error bars represent 95% confidence intervals around the effect estimate. Asterisks represent significantly nonzero marker effects at the p < 0.01 threshold.

**Figure 5.**
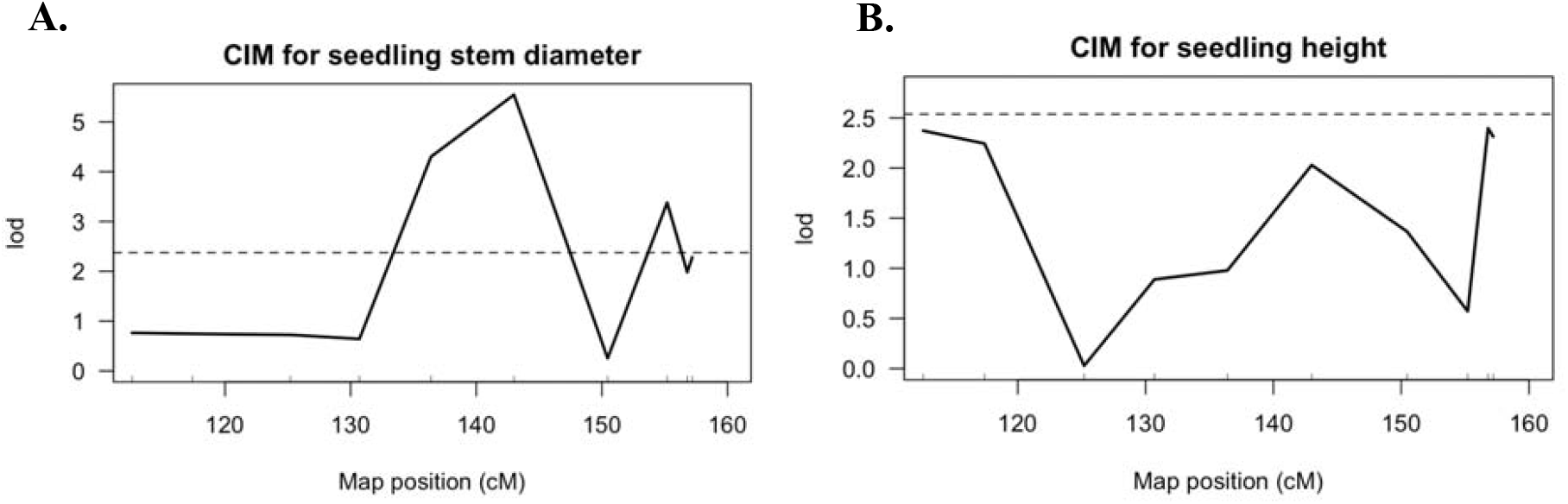
Composite interval mapping (CIM) results for A) seedling stem diameter and B) plant height at 4 MAP. The 10 C1GI markers are represented by the x-axis ticks according to their genetic position in cM on the distal end of chromosome 1. The y-axis gives the LOD score.

Among the clonal trials, stem diameter was also genetically correlated with vigor rating (τ = 0.18, p = 2.7x10^-8^) and plant height (τ = 0.42, p < 2.2x10^-16^). The estimated narrow-sense heritability was 0.45 for stem diameter and 0.70 for plant height (Table 2). The MLM method identified significant but small (∼0.69 mm) negative additive effects of C1GI markers 1 and 2 on clonal stem diameter (Fig. 6A), though these effects were not statistically significant with the CIM method (LOD = 0.69 and 0.02 respectively, Fig. 7A). Instead, with CIM, C1GI marker 8 had a significant LOD peak with small positive effects for both stem diameter (estimated additive effect = 0.22; dominance effect = 0.03 Fig. 7A) and vigor rating (estimated additive effect = 0.16; dominance effect = 0.27; Fig. 7B). There were no significant associations with clonal plant height using either the MLM or CIM methods (Fig. 6A, Fig. 7C).

**Figure 6.**
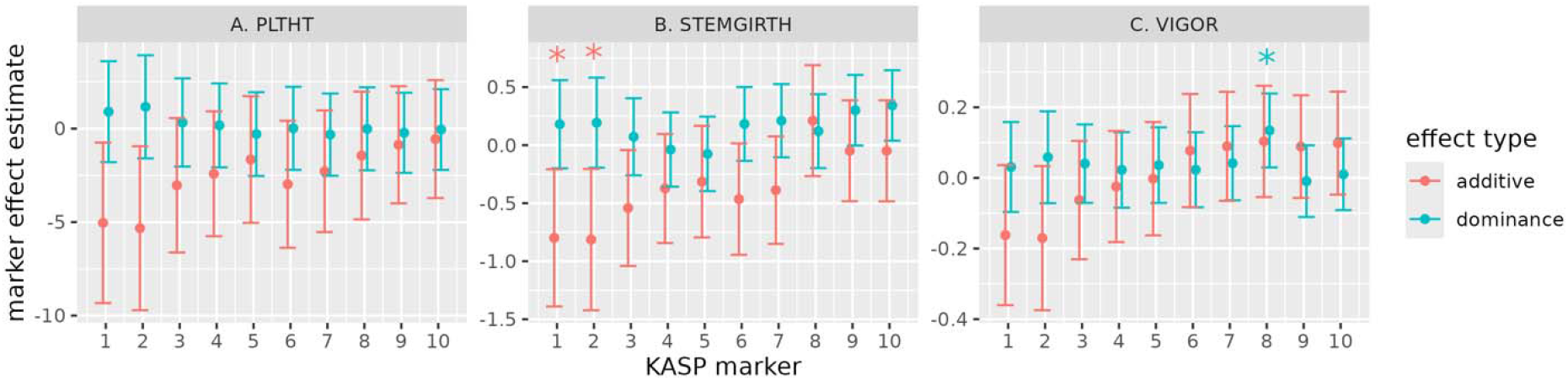
MLM estimates of additive (red) and dominance (blue) effects of *M. glaziovii* introgression markers on clonal vigor traits: A) plant height, B) stem diameter, and C) vigor rating. Error bars represent 95% confidence intervals around the effect estimate. Asterisks represent significantly nonzero marker effects at the p < 0.01 threshold from Wald tests.

**Figure 7.**
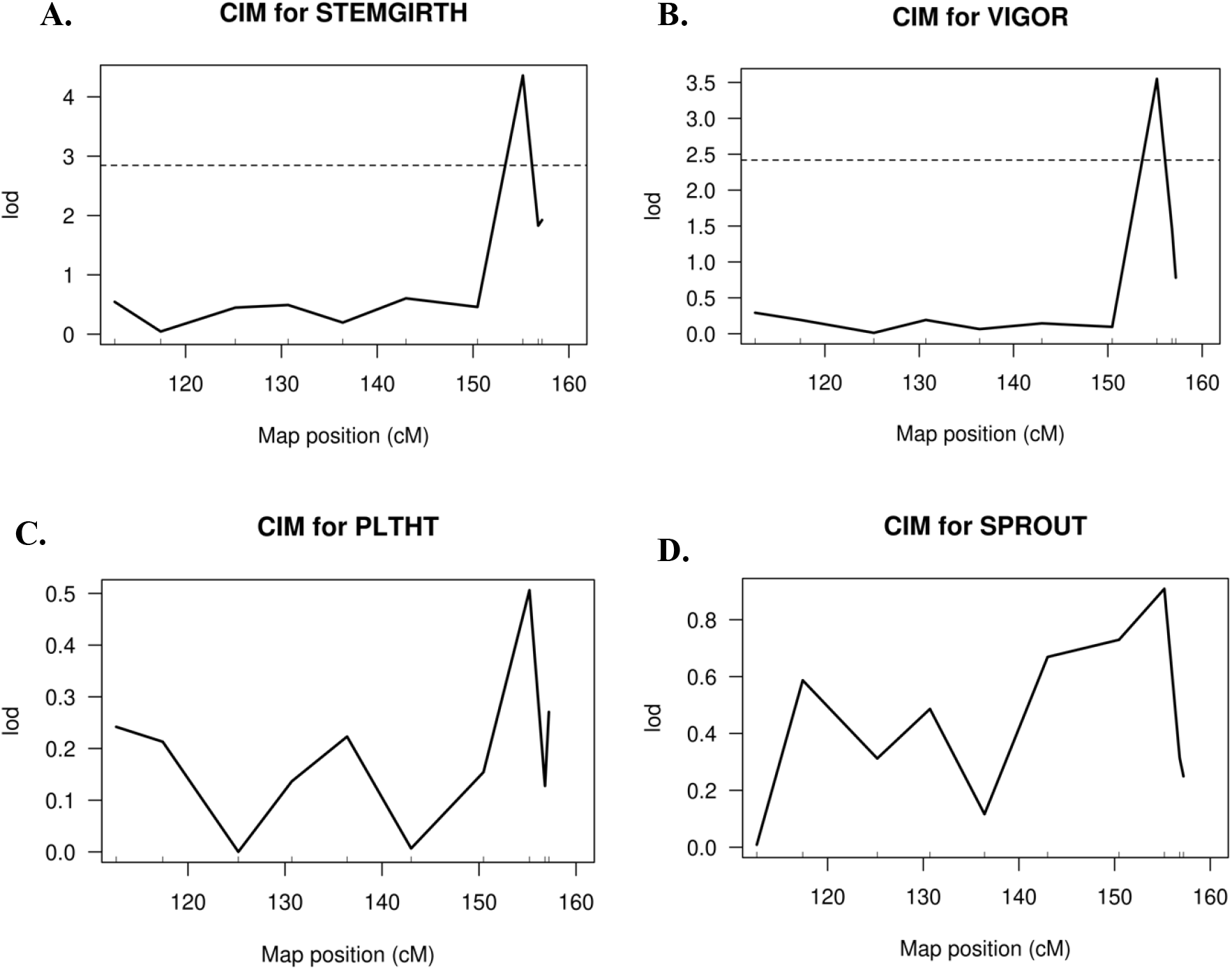
Composite interval mapping (CIM) results for clonal vigor traits: A) stem diameter, B) vigor score, C) plant height, and D) stem sprouting percentage. The 10 C1GI markers are represented by the x-axis ticks according to their genetic position in cM on the distal end of chromosome 1. The y-axis gives the LOD score.

**Figure 8.**
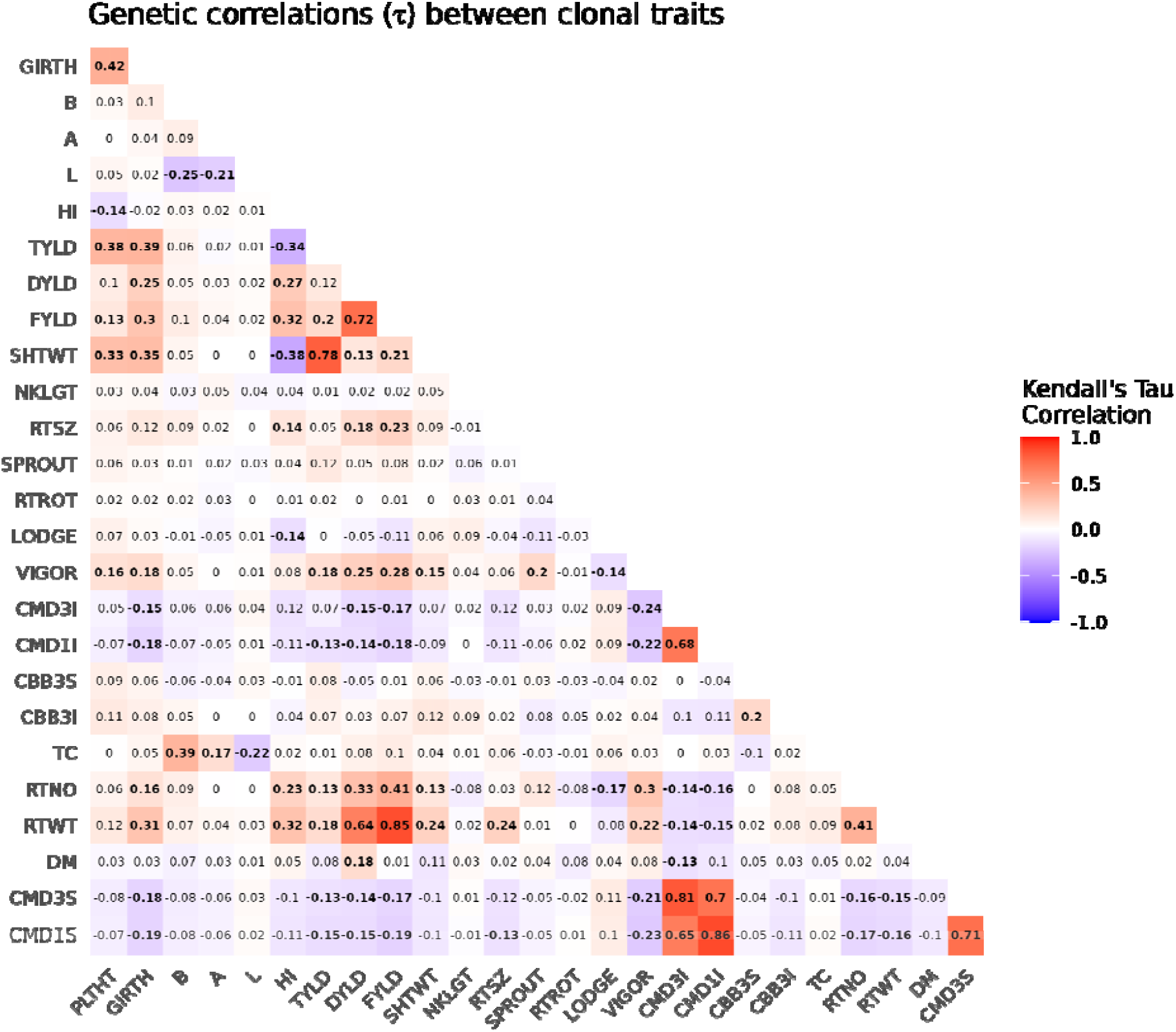
Genetic correlations between agronomic and clonal vigor traits. Values in each cell are pairwise Kendall’s Tau correlation values between de-regressed EBVs, bolded where statistically significant at the Bonferroni-corrected p < 0.00015 threshold.

### Agronomic trait association analyses

Across all agronomic traits listed in Table 2, the only significant C1GI marker/trait associations identified with the MLM method were for rotted root proportion, root neck length, and A chromameter value (Fig. 9). However, these marker/trait associations were not statistically significant in the CIM method accounting for multiple markers. With CIM, there was a marginally significant association between C1GI marker 8 and fresh root yield (additive effect = 0.42, dominance effect = 0.42; Fig. 10A).

**Figure 9.**
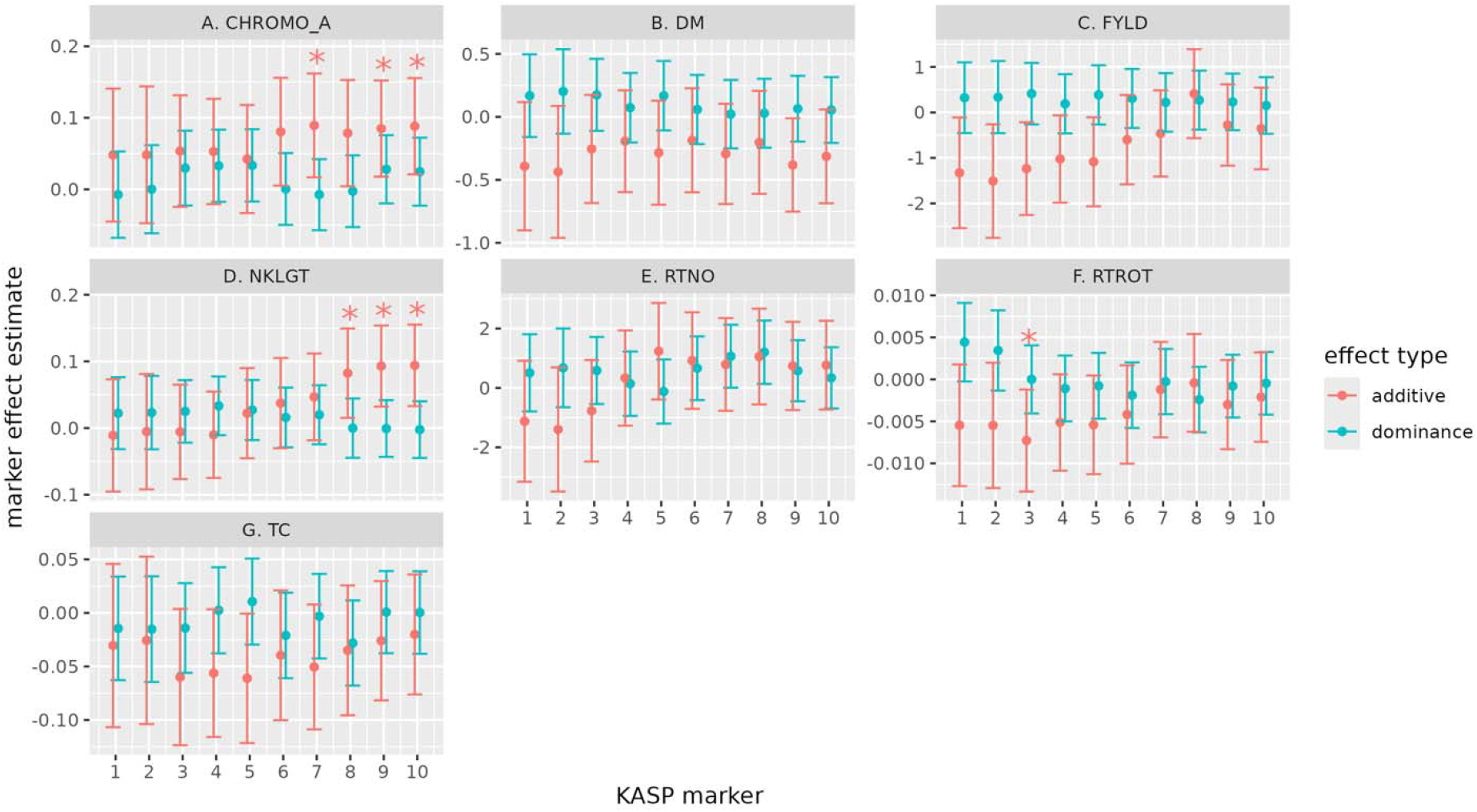
MLM estimates of additive (red) and dominance (blue) effects of C1GI markers on selected agronomic traits evaluated in clonal trials: A) chromameter a* value, B) dry matter content, C) fresh root yield, D) root neck length, E) root number, F) rotted root proportion, G) total carotenoids. Error bars represent 95% confidence intervals around the effect estimate. Asterisks represent significantly nonzero marker effects at the p < 0.01 threshold from Wald tests.

**Figure 10.**
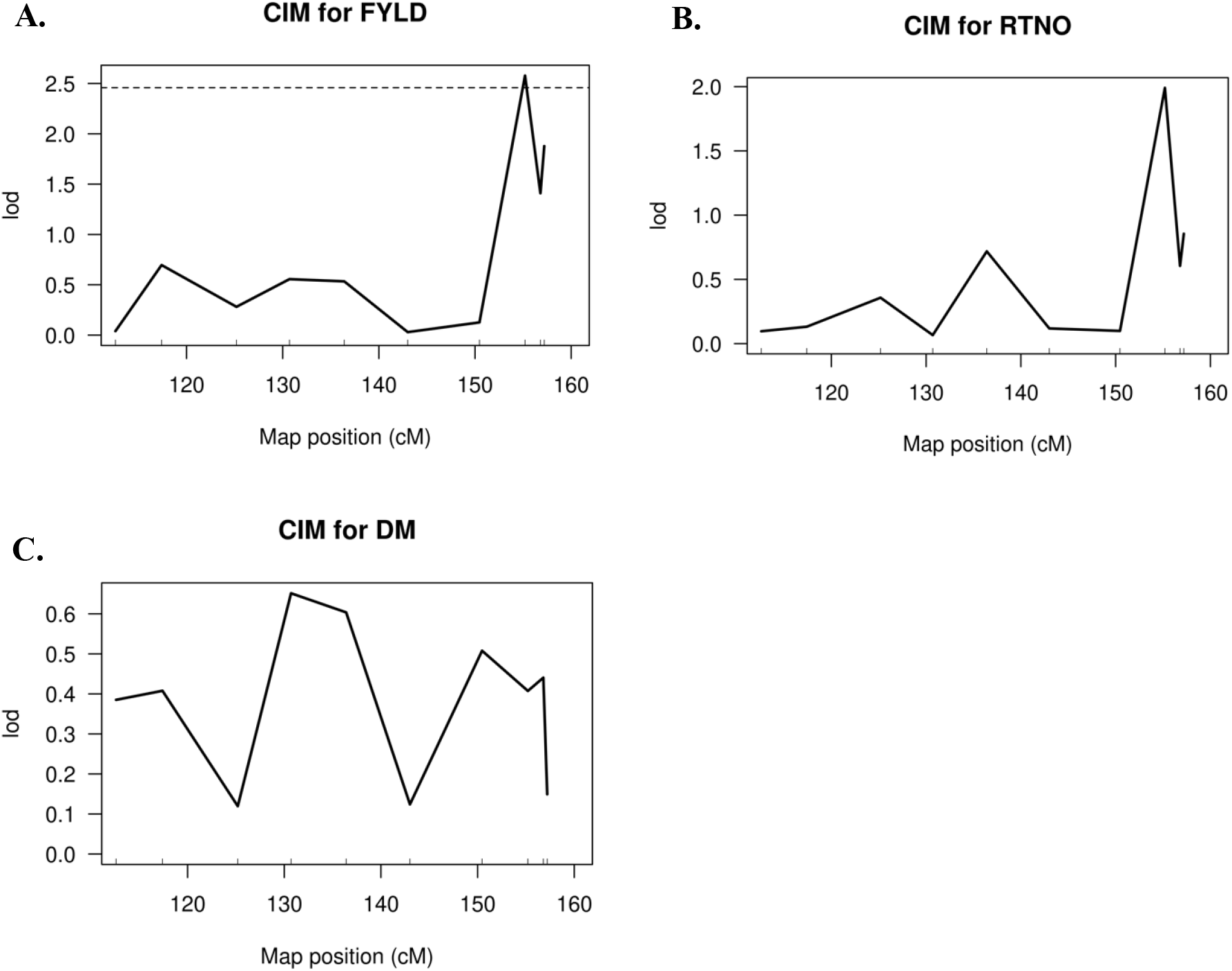
Composite interval mapping results for clonal yield traits: A) fresh root yield, B) root number, C) dry matter content. The 10 C1GI markers are represented by the x-axis ticks according to their genetic position in cM on the distal end of chromosome 1. The y-axis gives the LOD score.

Unexpectedly, there were no significant associations with DM or root number, the main traits that have been linked to the C1GI (Wolfe et al., 2019). DM and root number had moderate heritabilities of 0.41 and 0.21, respectively. Non-significant CIM results for the remaining agronomic traits are shown in Fig. S2.

Clonal stem diameter was positively genetically correlated with several yield measures: root weight (τ = 0.31, p < 2.2x10^-16^), root number (τ = 0.16, p = 8.8x10^-8^), root size (τ = 0.12, p = 0.0005), fresh root yield (τ = 0.31, p < 2.2x10^-16^), and dry root yield (τ = 0.25, p = 1.6x10^-14^), but not DM (τ =-0.03, p = 0.35).

### Genome alignment

The two *M. glaziovii* haplotype assemblies were aligned to the *M. esculenta* reference genome v8.1, which lacks the C1GI. Only insertions/deletions were detected in the C1GI region (Fig. 11). There were no apparent large inversions or duplications between *M. glaziovii* and *M. esculenta* chromosome 1 in the introgression region (Fig. 12). The largest structural variant in the C1GI region was an approximately 271kb deletion at 38.78 Mbp (Fig. 12). The full genome alignment is shown in Fig. S3 and the distribution of genome-wide structural variants called with SVIM-asm is shown in Fig. S4.

**Figure 11.**
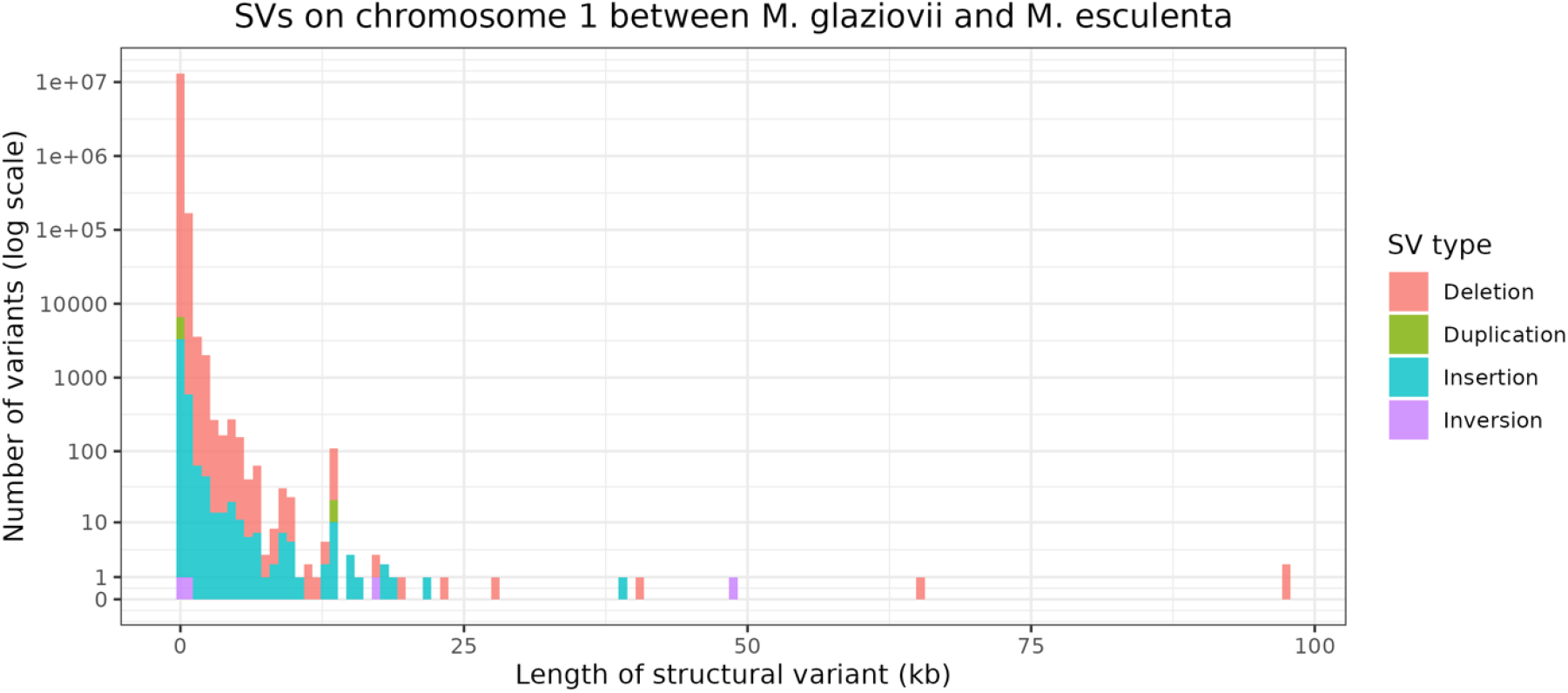
Distribution of structural variant (SV) sizes between *M. esculenta* and *M. glaziovii* genomes on chromosome 1. The x-axis represents the length of the structural variant in kilobases and the y-axis represents the number of variants on a log-transformed scale. SVs were called by SVIM-asm.

**Figure 12.**
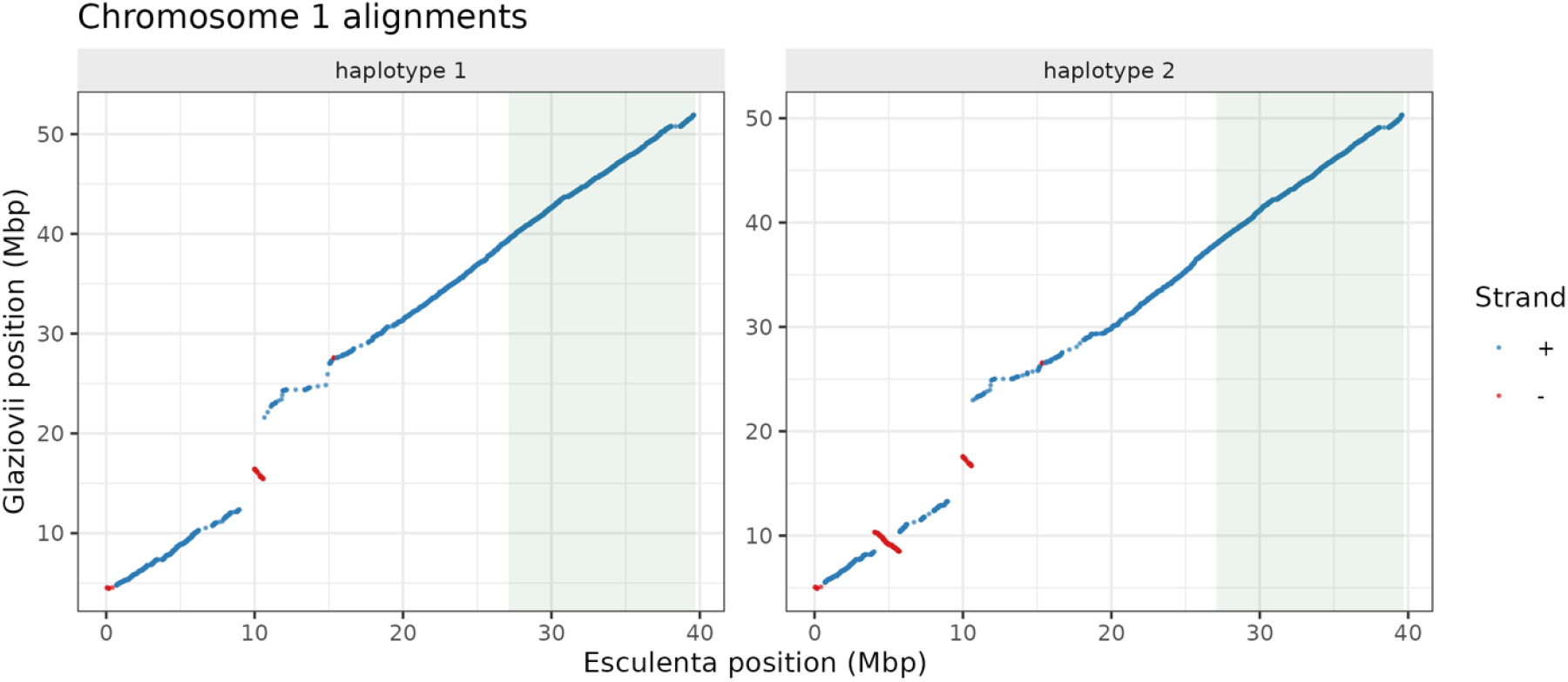
Alignment of *M. glaziovii* assembly v1 haplotype 1 (left) and haplotype 2 (right) to the *M. esculenta* reference genome v8.1 chromosome 1 show strong collinearity in the C1GI region. Dots represent physical positions of genes used as alignment anchors in the two genomes. The C1G1 region is highlighted in green.

### Enrichment of deleterious alleles

Across the 39,390 SNPs in the C1GI region, 1,602 were putative deleterious alleles predicted by Long et al. (2023) compared to 1,516 on average across 1,000 permutations of the same number of randomly selected sites in the genome. This difference corresponded to a p-value of 0.0096, indicating the C1GI is significantly enriched for deleterious alleles. The density of putative deleterious alleles across the genome is shown in Fig. 13.

**Figure 13.**
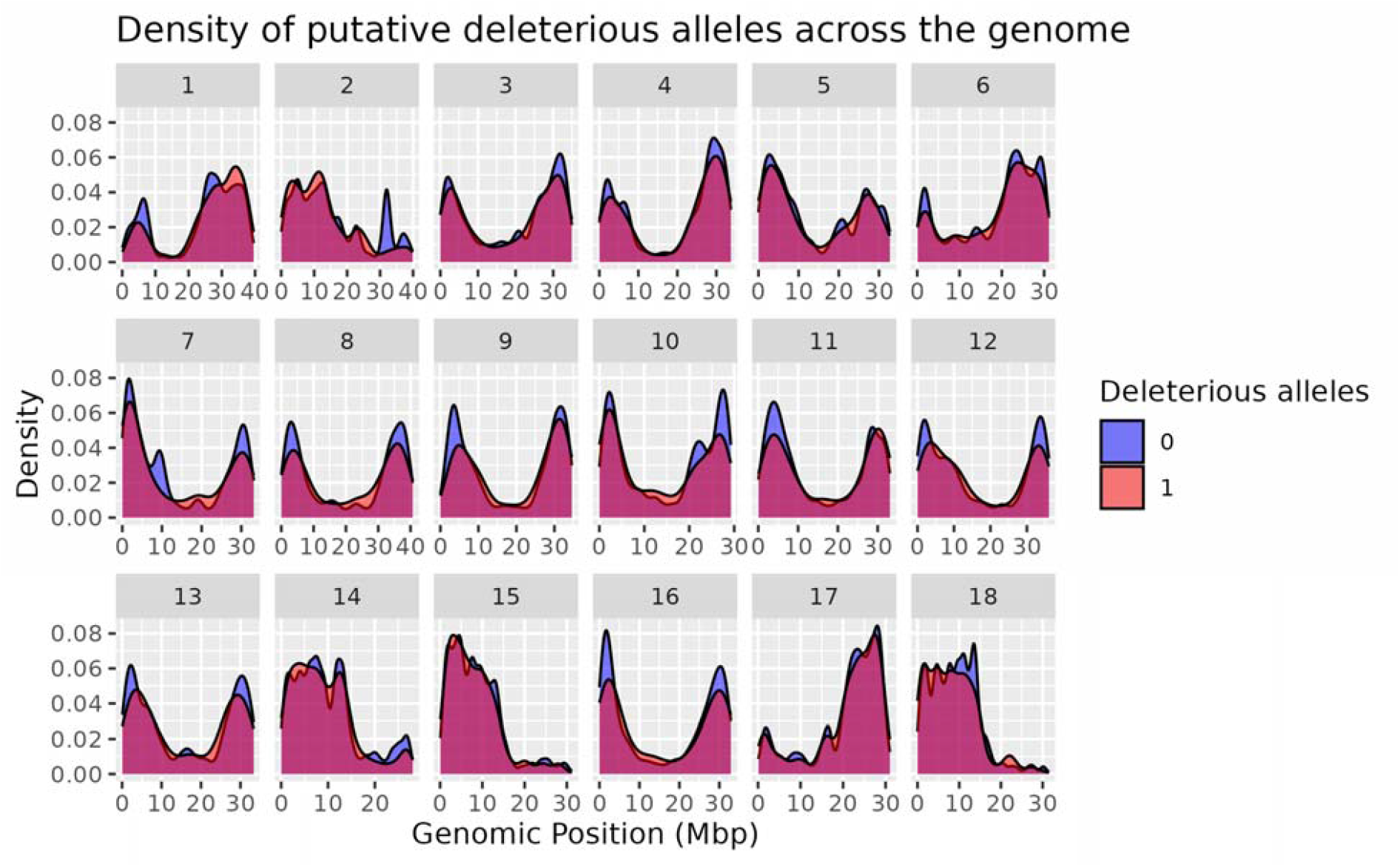
Density of putative deleterious alleles across the *M. esculenta* genome. Blue color (0) represents a non-deleterious SNP and red color (1) represents a deleterious SNP classified by Long et al. (2023) based on evolutionary conservation across the Euphorbiaceae family.

## 4 DISCUSSION

With the design of *glaziovii* allele specific KASP markers, a mapping population of 453 individuals with increased recombinations in the C1GI region was generated. Although QTL for DM and root number were previously identified in the C1GI (Wolfe et al., 2019), neither of these QTL were detected in this mapping population, despite the moderate heritability of DM estimated here (h^2^ = 0.41). One possible explanation is that there are multiple *glaziovii* haplotypes, and the one harboring the QTL was not captured in this population. However, given that the parents for this population were identified from a crossing nursery of parents selected in the genomic selection breeding pipeline, it is unlikely that the dry matter QTL would have been absent among them.

Alternatively, the QTL effect sizes could be too small to be detectable in this population of 453 compared to the population of over 4600 in the Wolfe et al. (2019) GWAS. *Glaziovii* introgression diagnostic markers contributed 0.05 to the heritability for DM and 0.01 to root number heritability in Wolfe et al. (2019). While associations with root number were not statistically significant (Fig. 9E, Fig. 10B), root number is positively correlated with fresh root yield (Fig. 8). C1GI marker 8 was associated with a positive effect on clonal fresh root yield (Fig. 10A). Therefore, the marginally-significant fresh root yield association supports the hypothesis that there was insufficient statistical power to detect the root number QTL in this population.

Finally, the original identification of the C1GI QTL may have been influenced by residual population structure. The presence of the C1GI has been considered an indicator of improved germplasm since it originated in the breeding programs, which have been selecting for DM. The Wolfe et al. (2019) GWAS accounted for relatedness with a kinship matrix constructed from non-*glaziovii* alleles, but it is possible that the genome-wide kinship relationships did not well capture the chromosome 1 structure. To test the hypothesis that the DM QTL is spurious, a stratified GWAS approach on subsets of the Wolfe et al. (2019) population could be used to determine whether the identification of the QTL is sensitive to the population subset.

The main C1GI marker-trait associations identified here were with aboveground vigor traits, both the breeder’s vigor ranking and stem diameter. There were similarities and differences between seedling and clonal vigor associations, which were analyzed separately since they reflect different biological processes (germination from seed versus clonal propagation). Both seedling and clonal CIM showed a significant LOD peak at marker 8 for stem diameter (Fig. 5A, Fig. 7A), with no significant peaks for plant height. There was an additional peak at C1GI markers 5-6 unique to seedling stem diameter (Fig. 5A). These results suggest the C1GI carries multiple QTL affecting stem thickness and therefore vigor at different developmental stages.

The positive correlation between stem diameter and vigor score, as well as the co-location of their associated markers, supports that stem thickness contributes to the breeder’s assessment of vigor. Assessing stem thickness in future trials may clarify its role in early screening criteria. The strongest selection intensity is applied at the SN stage, when populations are large, but relatively little attention is given to early selection decisions. Many lines with poor vigor are screened out at the SN stage before they are genotyped or included in training datasets for genomic prediction models. Therefore, the effects of recessive deleterious alleles may not be captured in genomic selection decisions if their expression is limited to homozygotes that are eliminated during early screening due to reduced vigor. Future studies could investigate incorporating predictions of deleterious alleles, such as those identified based on evolutionary constraint by (Long et al., 2023) or DNA language models by Zhai et al. (2024), into genomic prediction strategies.

Other significant C1GI associations were with chromometer a, root neck length, and rotted root proportion (Fig. 9). Chromometer a* value, representing redness vs. greenness, is not a particularly relevant metric alone, as it would usually be interpreted in combination with chromometer b* value measuring yellowness to estimate total carotenoid content (Bessa de Carvalho et al., 2022). Since shorter root neck lengths are generally preferred (International Institute of Tropical Agriculture, 1990), the association of *glaziovii* alleles at the distal end of C1GI with longer root neck length are in an unfavorable direction. Rotted root proportion is agronomically important, depending on the environment, but the association with C1GI was very small and only identified with the MLM method (Fig. 9F). Therefore, none of these agronomic trait associations are particularly useful for breeding. Like previous studies, we did not identify any associations with disease resistance traits (Fig. S2), which was the original impetus for making the *glaziovii* x *esculenta* crosses. Therefore, original motivation for making the *glaziovii* x *esculenta* crosses is no longer the current reason the C1GI is increasing in frequency. It remains unclear why the C1GI is increasing in frequency through the genomic selection pipeline. If the DM QTL is confounded with population structure, genomic predictions may be influenced by the spurious association in the same way as GWAS can.

Both CIM and MLM QTL mapping methods were used in this study for their relative strengths in accounting for multiple QTL simultaneously and in accounting for relatedness with a genomic relationship matrix. The CIM method demonstrated the ability to distinguish separate marker additive effects in opposite directions (Fig. 5A), which were not detected in the MLM method as they likely canceled each other out (Fig. 6B). While the MLM approach allows more flexibility to fit random effect covariates, the CIM method applies a variable selection algorithm to account for multiple markers simultaneously and distinguish their effects. Variable selection in mixed linear models is possible through methods like stepwise regression, though this approach has been criticized (Smith, 2018). Since the two methods accounted for population structure in different ways, some differences in the association results were expected, though the presence and absence of QTL between methods (e.g. the fresh root yield QTL identified with CIM but not MLM) suggests their sensitivity to population structure.

A key result from this study was the identification of segregation distortion of the *glaziovii* haplotype, with significantly fewer homozygous introgressed lines than expected, even after accounting for the rate of pedigree errors. This segregation distortion likely contributes to the observation of very few homozygous *glaziovii* lines in advanced trials (Wolfe et al., 2019). While early selection for vigor may also be a contributors, since the negative effects of *glaziovii* markers on seedling stem diameter were less than 1 mm, lower vigor perceived by the breeder does not fully explain why homozygous introgressed lines are excluded from advanced trials. Segregation distortion suggests that the C1GI carries deleterious alleles that affect either pollination success, germination, or survival. The enrichment of putative deleterious alleles in the C1GI region supports this hypothesis. Phasing the putative deleterious alleles in future studies would confirm whether they are enriched on the *glaziovii* haplotype relative to the *esculenta* haplotype in the C1GI region.

There were no apparent major chromosomal rearrangements that would explain the suppressed recombination in the C1GI region; the *glaziovii* haplotypes both showed strong synteny with the *esculenta* reference (Fig. 12). Suppressed recombination may instead be due to general sequence divergence between the *glaziovii* and *esculenta* haplotypes, which has been shown to affect crossover frequency in model organisms including *Arabidopsis* (Surtees et al., 2004; Li et al., 2006; Serra et al., 2018). However, there were many medium-sized insertions and deletions (Fig. 11), which may collectively contribute to degree of mismatch between the *esculenta* and *glaziovii* haplotypes and thus the suppressed recombination rates.

Given the observed segregation distortion, enrichment of putative deleterious alleles, and the lack of major beneficial QTL detected in the C1GI, recombining and partially or fully purging the introgression may be recommended. To facilitate this, a selection index weight could be placed on clones with recombinant C1GI segments. The C1GI KASP markers developed here may be useful for this purpose. They were added to a Amplicon-based SNP panel (3KCassCA) currently in development, which can be used to track the C1GI haplotype in the future.

Overall, this study reports a novel segregation distortion of the C1GI haplotype and a small but statistically significant association between the C1GI and poor plant vigor. Since the previously-reported QTL for dry matter content and root number could not be replicated, the benefit of maintaining the C1GI in cassava breeding populations warrants further evaluation. These findings demonstrate the long-term effects of wide crosses with wild relatives across many generations (∼80 years of breeding) without attention to purging linkage drag. Large introgression blocks from wild relatives should be carefully examined, since they can create regions of suppressed recombination that hinder favorable allelic combinations.

## Abbreviations

C1GI: chromosome 1 *glaziovii* introgression
CBSD: cassava brown streak disease
CMD: cassava mosaic disease
GWAS: genome-wide association study
KASP: ‘Kompetitive’ allele-specific PCR (polymerase chain reaction)
QTL: quantitative trait locus/loci.

## Data Availability Statement

Raw trial data are available on Cassavabase (https://www.cassavabase.org/) under the following trial names: 20.GS.C5.SN.IB, 21.GS.C5.CE.1600.IB, 22.GS.C6.STUDENT2_SEREN.CE.500.IK, 23.GS.C6.SEREN.PYT.50.IB, 23.GS.C6.SEREN.PYT.50.IK. Imputed DArTSeqLD genotype data are available on Cassavabase. C1GI KASP marker genotypes and all scripts used for analysis are publicly available at https://github.com/serenvillwock/chr1glaziovii.

## Conflict of Interest Statement

The authors declare no conflicts of interest.

## Author Contributions

Seren S. Villwock: Conceptualization; Methodology; Investigation; Data curation; Formal analysis; Project administration; Writing – original draft; Writing – review & editing. Ismail Y. Rabbi: Project administration; Resources; Funding acquisition; Supervision; Writing – review & editing. Andrew Smith Ikpan: Project administration; Investigation; Data curation; Resources; Supervision. Ogunpaimo Kayode: Investigation; Data curation. Nafiu Kehinde: Investigation; Data curation. Siraj Ismail Kayondo: Methodology. Marnin Wolfe: Conceptualization. Jean-Luc Jannink: Conceptualization; Methodology; Resources; Funding acquisition; Supervision; Writing – review & editing.

## Acknowledgements

We thank the Cornell University BRC Bioinformatics Core Facility (RRID:SCR_021757) for providing access to computational resources and Jeff Glaubitz for technical consultation. We are grateful to the Cassava Breeding Unit team at the International Institute of Tropical Agriculture, Ibadan, Nigeria, especially including Lynda Adaobi Nnadi, Peter Iluebby, Patrick Akpotuzor, Victor Monday Chukwuyem, Toye Ayankanmi, and Kehinde Nafiu for their assistance with tissue sampling and field trials; Prasad Peteti and Mahmud Kehinde Dhikrullah for their assistance with data curation; and Peter Kulakow, Peace Ikeanyi, and Richard Ofei for their administrative support. We are grateful to Bethany Econopouly, Todd Michael and team, and The Salk Institute for sharing the *Manihot glaziovii* genome assembly, to Mohamed El-Walid for technical consultation on AnchorWave, and to Evan Long for sharing the deleterious allele predictions and for technical consultation. The authors thank the UK’s Foreign, Commonwealth & Development Office (FCDO) and the Bill & Melinda Gates Foundation (Grant INV-007637 http://www.gatesfoundation.org) for their financial support. SSV is supported by the USDA National Institute of Food and Agriculture AFRI Predoctoral Fellowship project accession no. 1030847.

## Notes

### Competing Interest Statement

The authors have declared no competing interest.

## References

Abass, A.B., Towo, E., Mukuka, I., Ranaivoson, R., Tarawali, G., & Kanju, E. (2014). Growing Cassava: A Training Manual from Production to Postharvest. IITA, Ibadan, Nigeria.

Bates, D., Mächler, M., Bolker, B.M., & Walker, S.C. (2015). Fitting linear mixed-effects models using lme4. Journal of Statistical Software, 67. 10.18637/jss.v067.i01

Bessa de Carvalho, R.R., Marmolejo Cortes, D.F., Bandeira e Sousa, M., Alves de Oliveira, L., & Jorge de Oliveira, E. (2022). Image-based phenotyping of cassava roots for diversity studies and carotenoids prediction. PLOS ONE, 17, e0263326. 10.1371/journal.pone.0263326

Bredeson, J. V., Lyons, J.B., Prochnik, S.E., Wu, G.A., Ha, C.M., Edsinger-Gonzales, E., Grimwood, J., Schmutz, J., Rabbi, I.Y., Egesi, C., Nauluvula, P., Lebot, V., Ndunguru, J., Mkamilo, G., Bart, R.S., Setter, T.L., Gleadow, R.M., Kulakow, P., Ferguson, M.E., Rounsley, S., & Rokhsar, D.S. (2016). Sequencing wild and cultivated cassava and related species reveals extensive interspecific hybridization and genetic diversity. Nature Biotechnology, 34, 562–570. 10.1038/nbt.3535

Broman, K.W., Wu, H., Sen, S., & Churchill, G.A. (2003). R/qtl: QTL mapping in experimental crosses. Bioinformatics, 19, 889–890. 10.1093/bioinformatics/btg112

Browning, B.L., & Browning, S.R. (2016). Genotype Imputation with Millions of Reference Samples. American Journal of Human Genetics, 98, 116–126. 10.1016/j.ajhg.2015.11.020

Chan, A.W., Villwock, S.S., Williams, A.L., & Jannink, J.L. (2022). Sexual dimorphism and the effect of wild introgressions on recombination in cassava (Manihot esculenta Crantz) breeding germplasm. G3: Genes, Genomes, Genetics, 12. 10.1093/G3JOURNAL/JKAB372

Delame, M., Prado, E., Blanc, S., Robert-Siegwald, G., Schneider, C., Mestre, P., Rustenholz, C., & Merdinoglu, D. (2019). Introgression reshapes recombination distribution in grapevine interspecific hybrids. Theoretical and Applied Genetics, 132, 1073–1087. 10.1007/s00122-018-3260-x

Endelman, J.B. (2011). Ridge Regression and Other Kernels for Genomic Selection with R Package rrBLUP. The Plant Genome, 4, 250–255. 10.3835/plantgenome2011.08.0024

Fernandez-Pozo, N., Menda, N., Edwards, J.D., Saha, S., Tecle, I.Y., Strickler, S.R., Bombarely, A., Fisher-York, T., Pujar, A., Foerster, H., Yan, A., & Mueller, L.A. (2015). The Sol Genomics Network (SGN)-from genotype to phenotype to breeding. Nucleic Acids Research, 43, D1036–D1041. 10.1093/nar/gku1195

Fukuda, W.M.G., Guevara, C.L., Kawuki, R., & Ferguson, M.E. (2010). Selected morphological and agronomic descriptors for the characterization of cassava

Gao, X., Starmer, J., & Martin, E.R. (2008). A multiple testing correction method for genetic association studies using correlated single nucleotide polymorphisms. Genetic Epidemiology, 32, 361–369. 10.1002/gepi.20310

Garrick, D.J., Taylor, J.F., & Fernando, R.L. (2009). Deregressing estimated breeding values and weighting information for genomic regression analyses. Genetics Selection Evolution, 41, 1–8. 10.1186/1297-9686-41-55

Gemenet, D.C., da Silva Pereira, G., De Boeck, B., Wood, J.C., Mollinari, M., Olukolu, B.A., Diaz, F., Mosquera, V., Ssali, R.T., David, M., Kitavi, M.N., Burgos, G., Felde, T. Zum, Ghislain, M., Carey, E., Swanckaert, J., Coin, L.J.M., Fei, Z., Hamilton, J.P., Yada, B., Yencho, G.C., Zeng, Z.B., Mwanga, R.O.M., Khan, A., Gruneberg, W.J., & Buell, C.R. (2020). Quantitative trait loci and differential gene expression analyses reveal the genetic basis for negatively associated β-carotene and starch content in hexaploid sweetpotato [Ipomoea batatas (L.) Lam.]. Theoretical and Applied Genetics, 133, 23–36. 10.1007/s00122-019-03437-7

Haley, C.S., & Knott, S.A. (1992). A simple regression method for mapping quantitative trait loci in line crosses using flanking markers. Heredity, 69, 315–324. 10.1038/hdy.1992.131

Heller, D., & Vingron, M. (2020). SVIM-asm: Structural variant detection from haploid and diploid genome assemblies. Bioinformatics, 36, 5519–5521. 10.1093/bioinformatics/btaa1034

Hillocks, R.J., & Jennings, D.L. (2003). Cassava brown streak disease: A review of present knowledge and research needs. International Journal of Pest Management, 49, 225–234. 10.1080/0967087031000101061

Holland, J.B., Nyquist, W.E., & CervantesLMartínez, C.T. (2002). Estimating and Interpreting Heritability for Plant Breeding: An Update. Plant Breeding Reviews (pp. 9–112). Wiley.

International Institute of Tropical Agriculture. (1990). Cassava in Tropical Africa: A Reference Manual. In Chayce Publication Services (Ed.), Ibadan, Nigeria.

Lawrence, E.J., Griffin, C.H., & Henderson, I.R. (2017). Modification of meiotic recombination by natural variation in plants 68, 5471–5483. 10.1093/jxb/erx306

Li, H., Berent, E., Hadjipanteli, S., Galey, M., Muhammad-Lahbabi, N., Miller, D.E., & Crown, K.N. (2023). Heterozygous inversion breakpoints suppress meiotic crossovers by altering recombination repair outcomes. PLoS Genetics, 19. 10.1371/journal.pgen.1010702

Li, L., Jean, M., & Belzile, F. (2006). The impact of sequence divergence and DNA mismatch repair on homeologous recombination in Arabidopsis. The Plant Journal, 45, 908–916. 10.1111/j.1365-313X.2006.02657.x

Long, E.M., Romay, M.C., Ramstein, G., Buckler, E.S., & Robbins, K.R. (2023). Utilizing evolutionary conservation to detect deleterious mutations and improve genomic prediction in cassava. Frontiers in Plant Science, 13. 10.3389/fpls.2022.1041925

R Core Team. (2021). R: A Language and Environment for Statistical Computing

Rabbi, I.Y., Udoh, L.I., Wolfe, M., Parkes, E.Y., Gedil, M.A., Dixon, A., Ramu, P., Jannink, J., & Kulakow, P. (2017). GenomeLWide Association Mapping of Correlated Traits in Cassava: Dry Matter and Total Carotenoid Content. The Plant Genome, 10, 1–14. 10.3835/plantgenome2016.09.0094

Ramu, P., Esuma, W., Kawuki, R., Rabbi, I.Y., Egesi, C., Bredeson, J. V., Bart, R.S., Verma, J., Buckler, E.S., & Lu, F. (2017)(a). Cassava haplotype map highlights fixation of deleterious mutations during clonal propagation. Nature Genetics, 49, 959–963. 10.1038/ng.3845

Ramu, P., Esuma, W., Kawuki, R., Rabbi, I.Y., Egesi, C., Bredeson, J. V., Bart, R.S., Verma, J., Buckler, E.S., & Lu, F. (2017)(b). Cassava haplotype map highlights fixation of deleterious mutations during clonal propagation. Nature Genetics, 49, 959–963. 10.1038/ng.3845

Rowan, B.A., Heavens, D., Feuerborn, T.R., Tock, A.J., Henderson, I.R., & Weigel, D. (2019). An Ultra High-Density Arabidopsis thaliana Crossover. Genetics, 213, 771–787

Schmidt, P., Hartung, J., Rath, J., & Piepho, H.P. (2019). Estimating broad-sense heritability with unbalanced data from agricultural cultivar trials. Crop Science, 59, 525–536. 10.2135/cropsci2018.06.0376

Serra, H., Choi, K., Zhao, X., Blackwell, A.R., Kim, J., & Henderson, I.R. (2018). Interhomolog polymorphism shapes meiotic crossover within the Arabidopsis RAC1 and RPP13 disease resistance genes. PLoS Genetics, 14. 10.1371/journal.pgen.1007843

Smith, G. (2018). Step away from stepwise. Journal of Big Data, 5. 10.1186/s40537-018-0143-6

Song, B., Marco-Sola, S., Moreto, M., Johnson, L., Buckler, E.S., & Stitzer, M.C. (2022). AnchorWave: Sensitive alignment of genomes with high sequence diversity, extensive structural polymorphism, and whole-genome duplication. PNAS, 119. 10.1073/pnas.2113075119

Surtees, J.A., Argueso, J.L., & Alani, E. (2004). Mismatch repair proteins: key regulators of genetic recombination. Cytogenetic and Genome Research, 107, 146–159. 10.1159/000080593

Tahirou, A., Bamire, A.S., Oparinde, A., & Akinola, A.A. (2015). Determinants of Adoption of Improved Cassava Varieties among Farming Households in Oyo, Benue, and Akwa Ibom States of Nigeria. HarverstPlus Working Paper, 22, 21

The VSNi Team. (2023). asreml: Fits Linear Mixed Models using REML

Whalen, A., Gorjanc, G., & Hickey, J.M. (2019). Parentage assignment with genotyping-by-sequencing data. Journal of Animal Breeding and Genetics, 136, 102–112. 10.1111/jbg.12370

Wheeler, B. (2019). AlgDesign: Algorithmic Experimental Design. R package version 1.2.0.

Wolfe, M. (2022). genomicMateSelectR: Genomic Mate Selection. R package version 0.2.0.

Wolfe, M.D., Bauchet, G.J., Chan, A.W., Lozano, R., Ramu, P., Egesi, C., Kawuk, R., Kulakow, P., Rabbi, I., & Jannink, J.L. (2019)(a). Historical introgressions from a wild relative of modern cassava improved important traits and may be under balancing selection. Genetics, 213, 1237–1253. 10.1534/genetics.119.302757

Wolfe, M.D., Bauchet, G.J., Chan, A.W., Lozano, R., Ramu, P., Egesi, C., Kawuk, R., Kulakow, P., Rabbi, I., & Jannink, J.L. (2019)(b). Historical introgressions from a wild relative of modern cassava improved important traits and may be under balancing selection. Genetics, 213, 1237–1253. 10.1534/genetics.119.302757

Zeng, Z.B. (1994). Precision mapping of quantitative trait loci. Genetics, 136, 1457–1468. 10.1093/genetics/136.4.1457

Zhai, J., Gokaslan, A., Schiff, Y., Berthel, A., Liu, Z.-Y., Lai, W.-Y., Miller, Z.R., Scheben, A., Stitzer, M.C., Romay, C., Buckler, E.S., & Kuleshov, V. (2024). Cross-species modeling of plant genomes at single nucleotide resolution using a pre-trained DNA language model. bioRxiv [preprint],. 10.1101/2024.06.04.596709

