## Supplemental Information for "Introgression from the wild relative *Manihot glaziovii* on cassava (*M. esculenta*) chromosome 1 exhibits segregation distortion and no direct effect on dry matter"

**Supplementary Information**

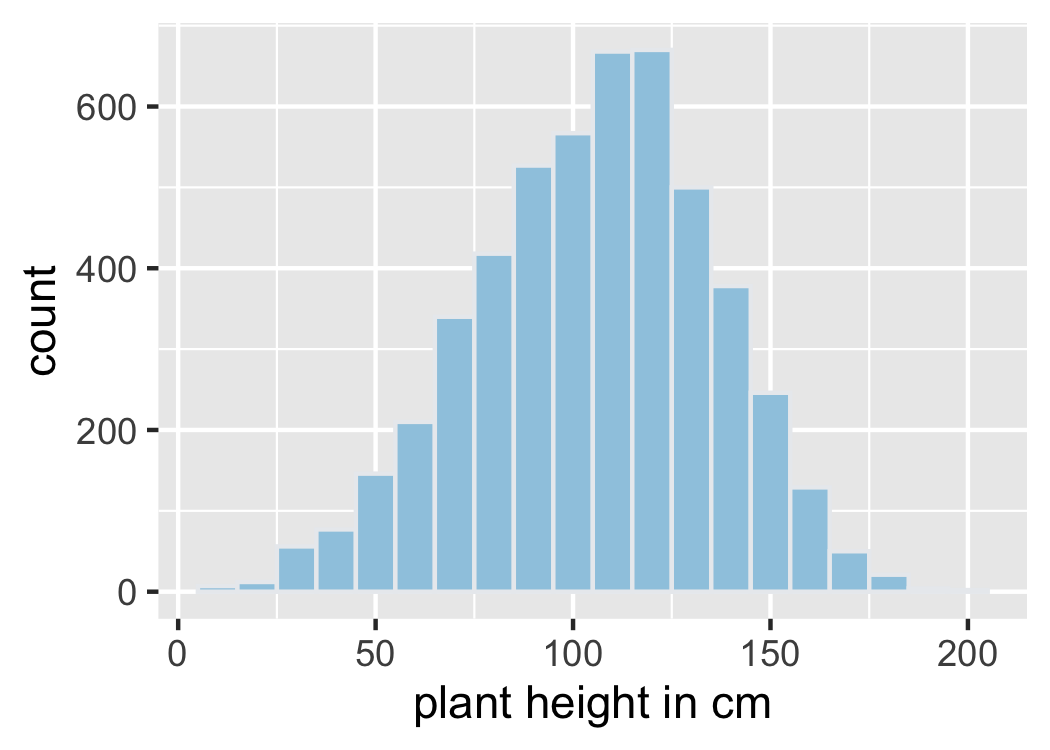

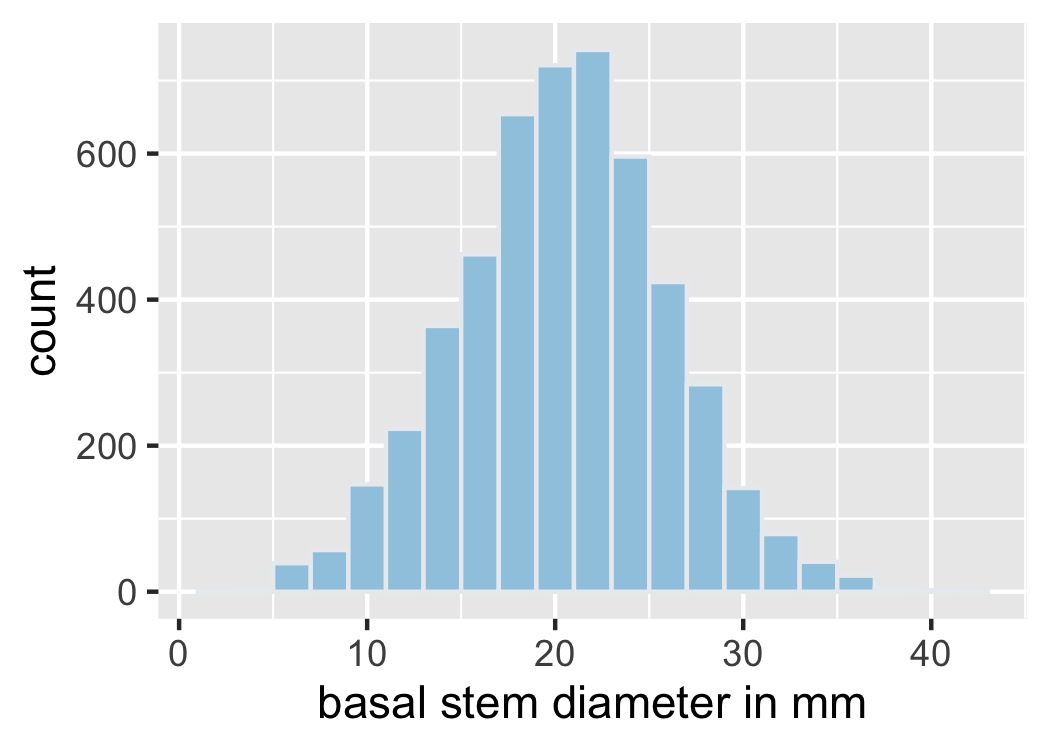

**Figure S1.** Distribution of vigor component traits stem diameter and plant height among 5000 seedlings at 4 MAP.

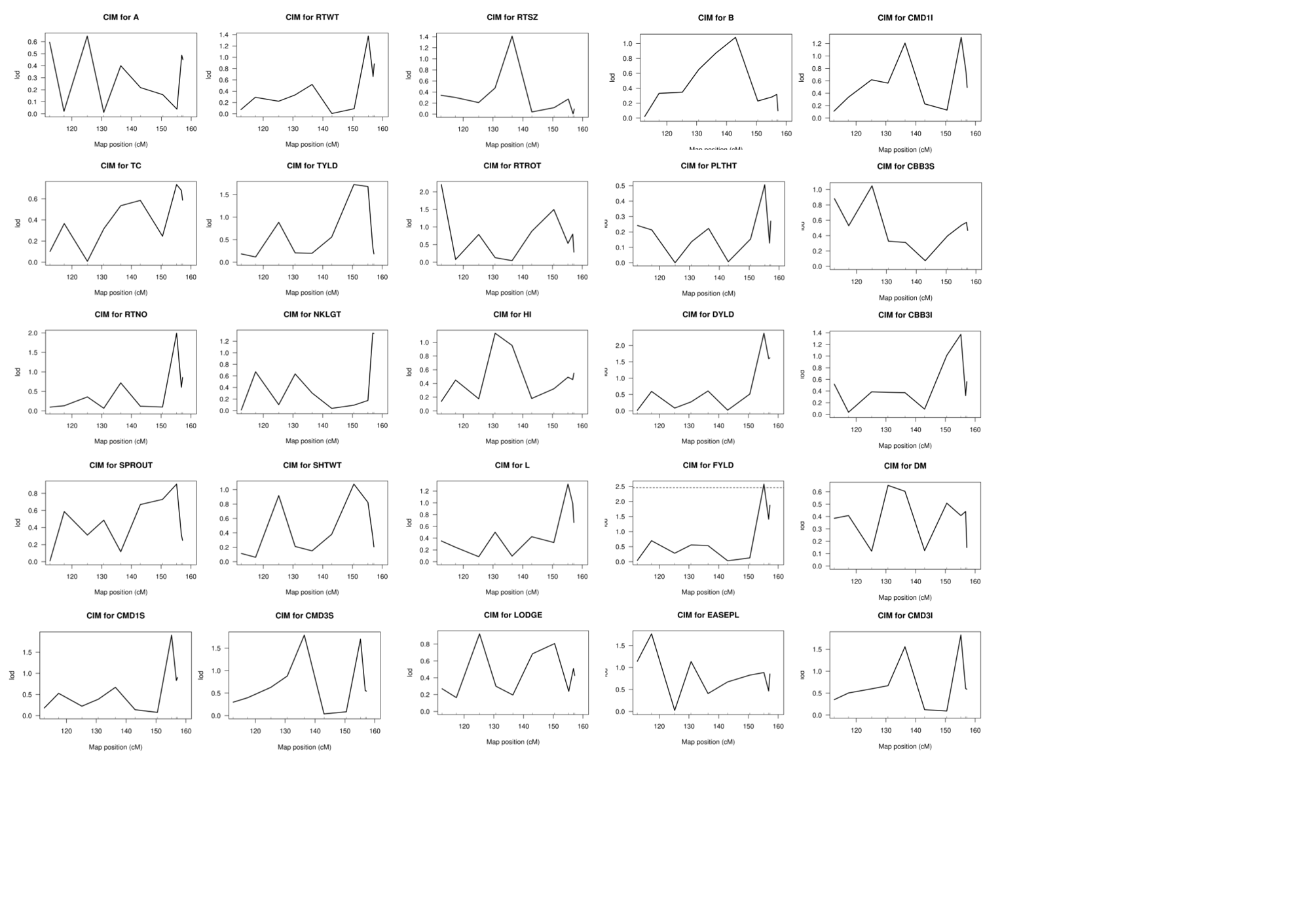

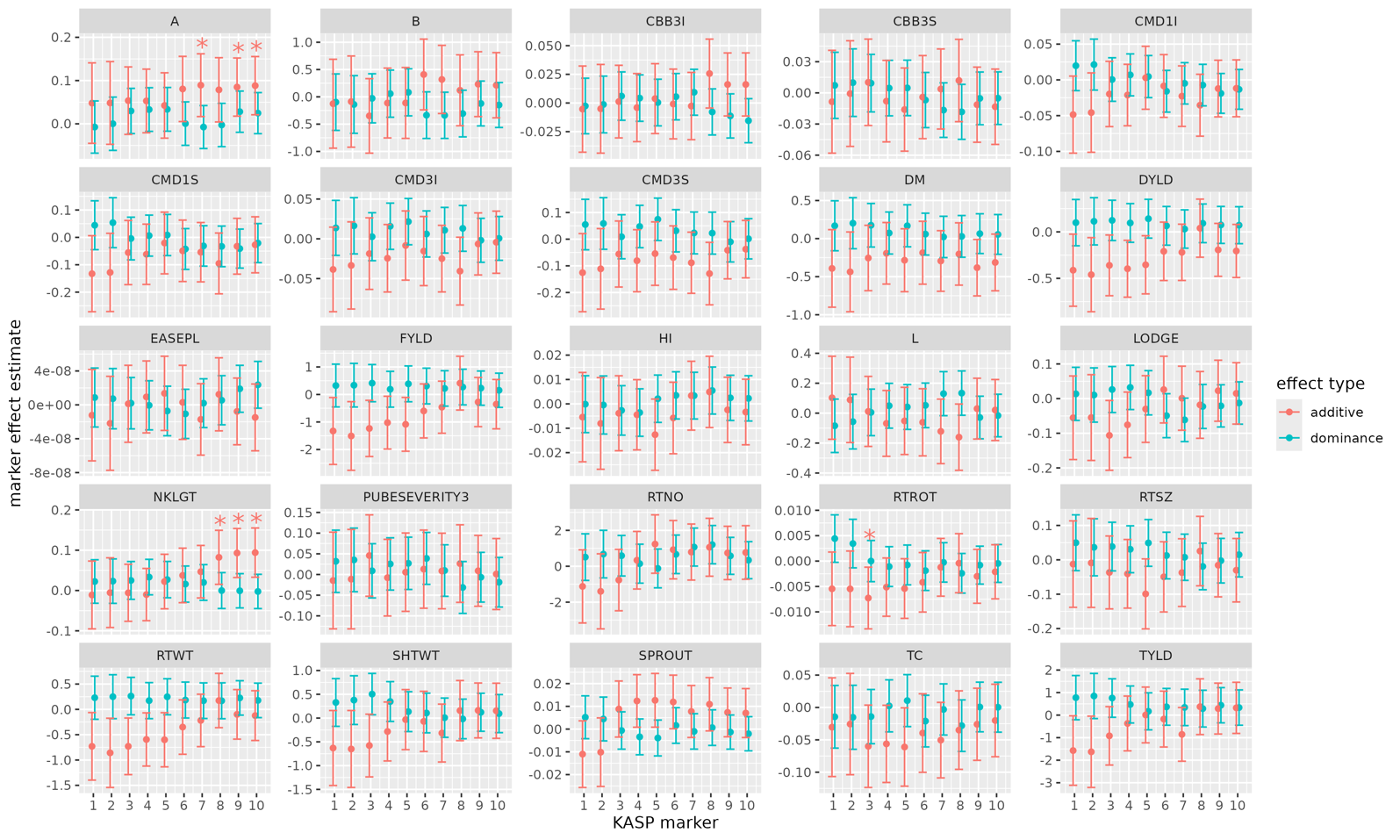

**Figure S2.** CIM (top panel) and MLM (bottom panel) association results for C1GI marker effect estimates on all agronomic traits measured in clonal trials.

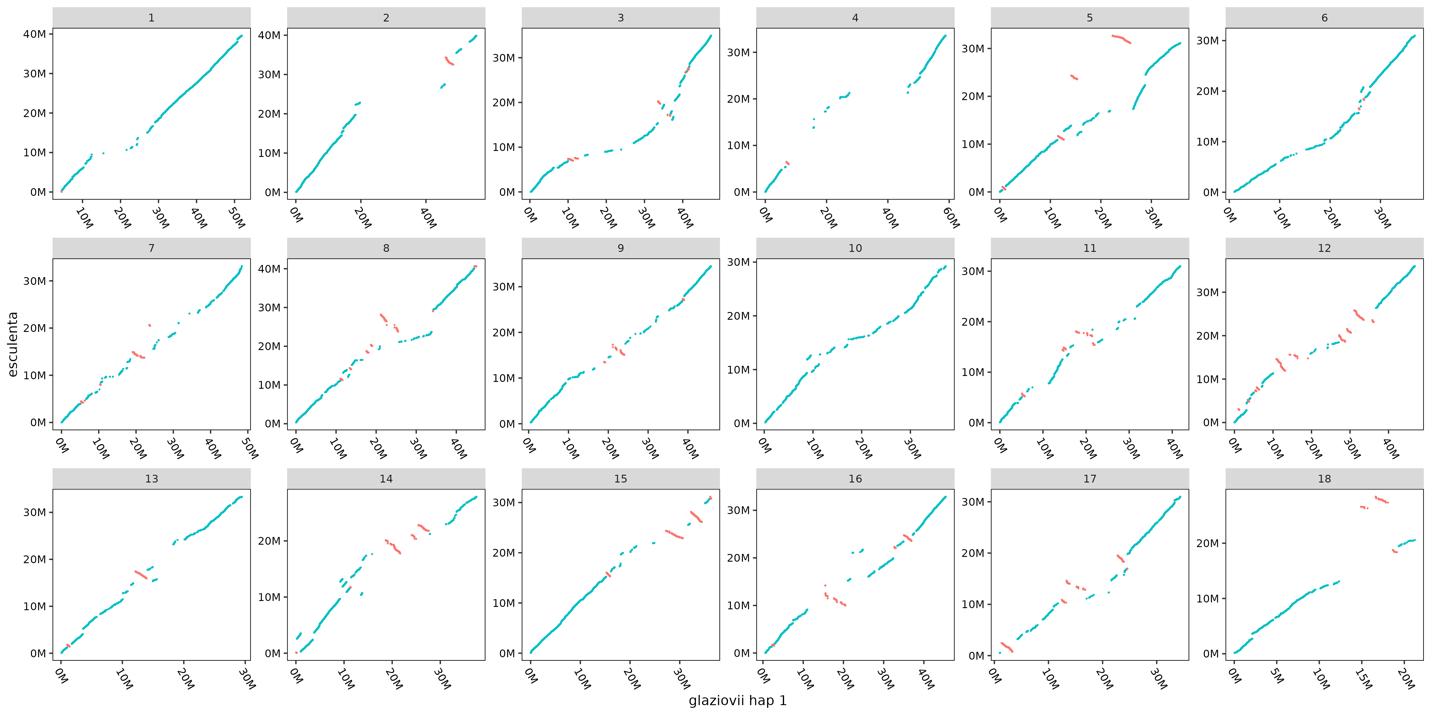

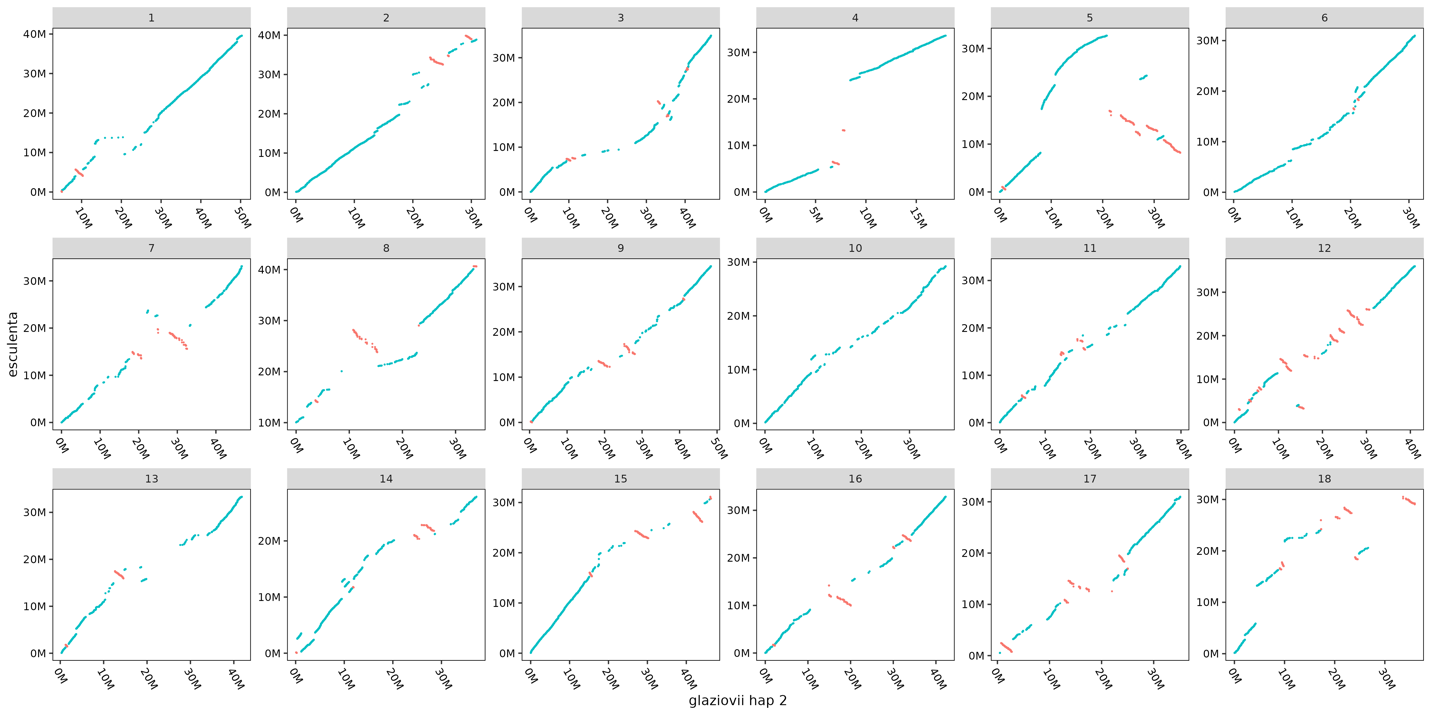

**Figure S3.** Genome-wide chromosome alignment between *M. esculenta* reference genome v8.1 and *M. glaziovii* haplotype 1 (top panel) and haplotype 2 (bottom panel) assemblies.

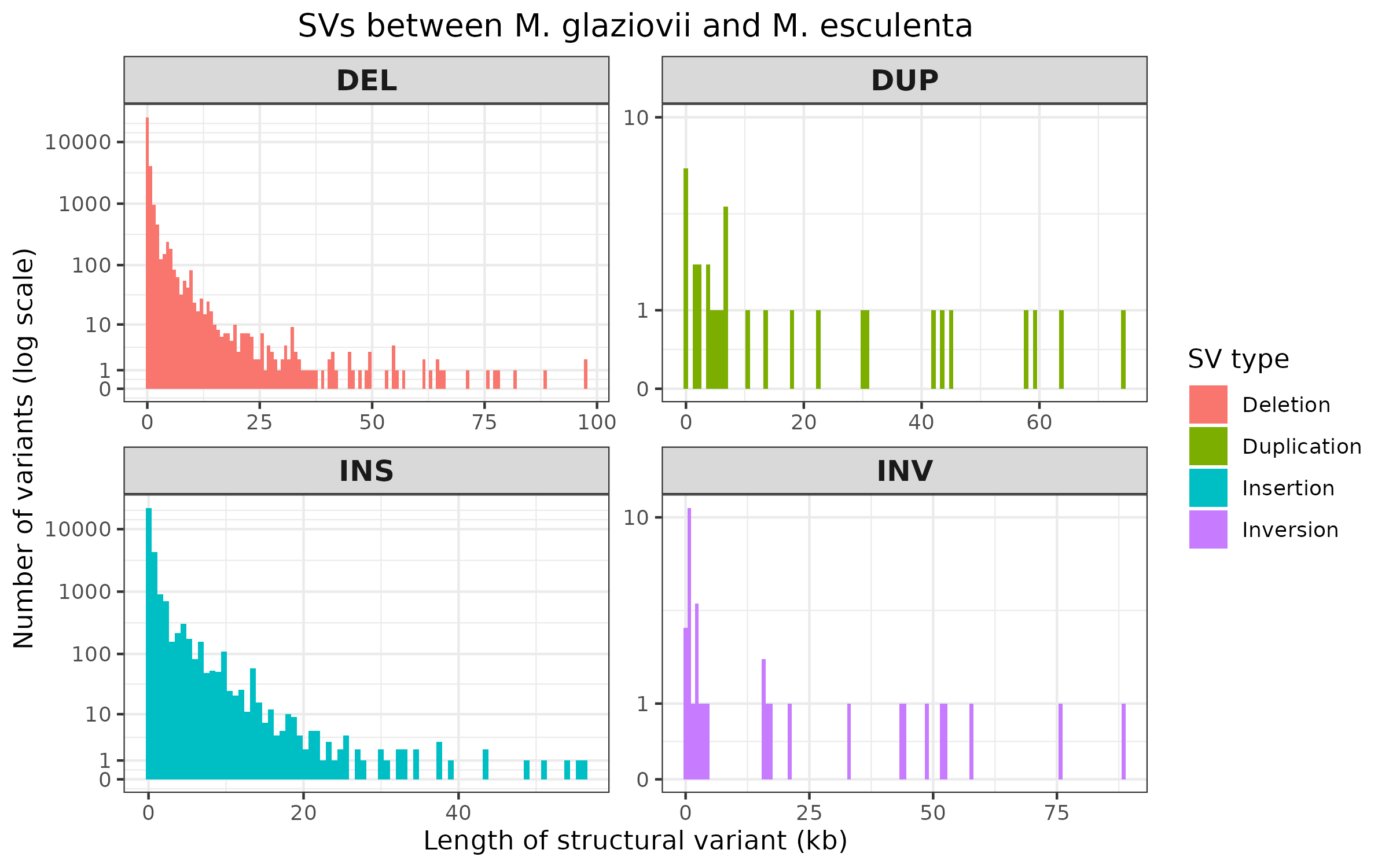

**Figure S4.** Distribution of structural variant (SV) types and lengths between *M. esculenta* reference genome v8.1 and the *M. glaziovii* haplotype 1 and 2 assemblies. SVs were called with SVIM-asm.
